# The *Phytophthora capsici* RxLR effector *CRISIS2* has roles in suppression of PTI and triggering cell death in host plant

**DOI:** 10.1101/2022.03.21.485173

**Authors:** Hyunggon Mang, Ye-Eun Seo, Hye-Young Lee, Haeun Kim, Hyelim Jeon, Xin Yan, Subin Lee, Sang A Park, Myung-Shin Kim, Cécile Segonzac, Doil Choi

## Abstract

Pathogen effectors can suppress various plant immune responses, suggesting that they have multiple targets in the host. To understand the mechanisms underlying plasma membrane-associated and effector-mediated immunity, we performed *Phytophthora <u>c</u>apsici* <u>R</u>xLR cell death <u>I</u>nducer <u>S</u>uppressing <u>I</u>mmune <u>S</u>ystem (CRISIS) screening. In *Nicotiana benthamiana*, the cell death induced by the RxLR effector CRISIS2 is inhibited by the irreversible plasma membrane H^+^-ATPase (PMA) activator fusicoccin. Biochemical and gene silencing analyses revealed that CRISIS2 physically and functionally associates with PMAs and induces host cell death independent of immune receptors. CRISIS2 induces apoplastic alkalization by suppressing PMA activity via its association with the C-terminal regulatory domain of PMA. *In planta* expression of CRISIS2 significantly enhanced the virulence of *P. capsici*, whereas host-induced gene silencing of CRISIS2 compromised disease symptom and biomass of *P. capsici*. Furthermore, co-immunoprecipitation assays revealed that CRISIS2 constitutively associates with BAK1, the co-receptor of pattern recognition receptors (PRRs). CRISIS2 interferes with the FLS2-BAK1 complex induced by flagellin perception and impairs downstream signaling from the PRR complex. Proteomics and gene silencing assays identified putative PRRs that negatively regulate the virulence of *P. capsici* in *N. benthamiana* as interactors of CRISIS2 and BAK1. Our study identified a novel RxLR effector playing multiple roles in the suppression of plant defense and induction of cell death to support the pathogen hemibiotrophic life cycle in the host plant.

## Introduction

Plants evolutionarily acquire an innate immune system to survive against invading pathogens. In the host cell, immunity is triggered by the perception of foreign components by extra- and intracellular immune receptors. The first layer of the immune system, known as pattern-triggered immunity (PTI), is activated by pathogen-associated molecular patterns (PAMPs) that are recognized by plasma membrane (PM)-localized pattern recognition receptors (PRRs) (Boller and He, 2009; Monaghan and Zipfel, 2012). Both eukaryotic and prokaryotic pathogens deliver effector proteins into the host cell to attenuate PTI at various levels. In the second layer of plant immunity, intracellular immune receptors termed, nucleotide binding leucine-rich repeat receptors (NLRs), recognize pathogen effectors in resistant hosts and result in effector-triggered immunity (ETI) (Maekawa et al., 2011). Generally, avirulent effector is perceived by corresponding NLR protein, and this perception triggers robust programmed cell death known as the hypersensitive response (HR). NLR proteins often require HSP90 (heat shock protein 90) which forms a complex including SGT1 (suppressor of the G2 allele of *skp1*) and RAR1 (required for Mla12 resistance) to induce HR, Thus the complex of HSP90-SGT1-RAR1 is considered as an important component of immune-chaperones in plants (Botër et al., 2007).

At the onset of pathogenic molecule perception, numerous cellular responses are regulated by PM-localized enzymes or ion channels (Boller and Felix, 2009). Plasma membrane H^+^-ATPases (PMAs) are primary pumps that build up an electrochemical gradient across the PM, which is essential for all living organisms (Kühlbrandt, 2004). During PTI, the activity of PMAs is dynamically regulated and therefore targeted by multiple pathogens. Various PAMPs, such as the fungal cell wall component chitin, a *Phytophthora megasperma* oligopeptide of 13 amino acids (Pep-13), or the bacterial flagellin active epitope flg22, induce ion fluxes and rapid alkalization of the medium in plant cell suspension cultures, likely through the inhibition of PMA activity (Felix et al., 1999; Felix et al., 1993; Nürnberger et al., 1994). Conversely, acidification of the apoplast often occurs during ETI. The Avr5 effector from *Cladosporium fulvum* is recognized by the corresponding NLR protein Cf5 and this results in the activation of PMAs followed by acidification of the extracellular medium in tomato cell suspension culture (Vera-Estrella et al., 1994). Moreover, the interaction between barley harboring the Mla3 protein and the avirulence effector AvrMla3 from *Blumeria graminis* f. sp. *hordei* (*Bgh*) also causes apoplastic acidification (Zhou et al., 2000). However, in other cases, the interaction between barley and *Bgh* mediated by different R (resistance) proteins induces apoplastic alkalization (Felle et al., 2004). Thus, disturbance of the electrochemical gradient across the PM appears to be a general feature of pathogen infection, but the molecular mechanism governing apoplastic acidification or alkalization is still unclear. Recently, several PM-associated coiled-coil (CC) domains of NLRs (CNLs) have been reported to inhibit host PMA activity, resulting in apoplastic alkalization leading to cell death (Lee et al., 2022). Although the apoplastic pH balance regulated by PMA activity plays an important role in diverse plant-pathogen interactions, how PMA activity is affected by various effectors remains to be elucidated.

The *Phytophthora* genus in the eukaryotic group of Oomycetes contains numerous devastating plant pathogens, such as *P. sojae*, *P. capsici*, or *P. infestans* (Tyler et al., 2006). Most of them are hemibiotrophic pathogens, presenting an initial biotrophic stage in which they take nutrients from living host cells followed by a necrotrophic stage during which the pathogen kills the host cells to take nutrients from the dead tissue (Koeck et al., 2011). Each *Phytophthora* species is armed with hundreds of effectors containing a conserved N-terminal Arg-any amino acid-Leu-Arg (RxLR) motif mediating their translocation into the host cell for manipulation of immune responses (Dou et al., 2008; Tyler et al., 2006; Whisson et al., 2007). In recent decades, many studies have focused on identifying the host targets of the RxLR effectors to understand the mechanisms of pathogenicity. For instance, the *P. infestans* effector AVRblb2 associates with the host papain-like protease C14 to prevent its secretion into the apoplast, resulting in enhanced susceptibility of host plants to *P. infestans* (Bozkurt et al., 2011). Another *P. infestans* RxLR effector, AVR3a, manipulates plant immunity by stabilizing the host E3 ligase CMPG1 (Bos et al., 2010). Recently, the *P. capsici* effector RxLR207 has been reported to manipulate host immunity by binding and degrading BPA1 (binding partner of ACD11) and four other BPLs (BPA1-like proteins), which are regulators of the reactive oxygen species (ROS)-mediated defense response (Li et al., 2019).

Several pathogen effectors are known to have multiple targets. As an example, *Pseudomonas syringae* AvrPto is localized to the PM of host cells, where it interacts with the intracellular Ser/Thr protein kinase Pto and, in concert with the NLR protein Prf, activates ETI in tomato (Mucyn et al., 2006). It has also been reported that AvrPto associates with the kinase domain of different Arabidopsis PRRs, including the receptor-like kinases (RLKs) FLS2 (Flagellin-Sensitive 2) and EFR (EF-Tu receptor), hence blocking PTI signaling through inhibition of the FLS2 or EFR kinase activity (Shan et al., 2008; Xiang et al., 2008). Recently several studies suggested that L-type lectin receptor-like kinases (LecRLKs), a plant-specific family of receptor kinase, could be potential immune receptors for *Phytophthora* genus, as they play an important role in plant immunity (Wang et al., 2014; Wang et al., 2015). However, the extent to which plant pathogen effectors interfere with defense-related RLKs at the PM is largely unknown.

To understand the mechanisms underlying PM-associated and effector-mediated plant innate immunity, we developed a series of multiomics and bioinformatics tools for screening the *P. capsici* RxLR effectors affecting the activity of PMAs and the associated cell death. We employed fusicoccin, an irreversible PMA activator, which inhibits the cell death triggered by PM-associated CNL (Lee et al., 2022). Here, we report that a novel *P. capsici* RxLR effector, CRISIS2, induces NLR-independent cell death by inhibiting PMA activity. CRISIS2 binding at the autoregulatory domain of NbPMA3 results in apoplastic alkalization and cell death. Interestingly, CRISIS2 also constitutively associates with the FLS2-BAK1 complex and interrupts the flg22-induced interaction between FLS2 and BAK1. By coimmunoprecipitation/tandem mass spectrometry (coIP/MS) assay, we further identified putative PRRs which are involved in basal defense against *P. capsici* and interact with CRISIS2 and NbBAK1. Collectively, we show that CRISIS2 has a dual role in the pathogenicity of *P. capsici* by inhibiting PTI through association with PRR complexes and by inducing host cell death through disturbance of the PMA activity in the host *N. benthamiana*.

## Results

### Screening of *Phytophthora <u>c</u>apsici* <u>R</u>xLR cell death <u>I</u>nducer <u>S</u>uppressing <u>I</u>mmune <u>S</u>ystem (CRISIS)

PMAs are targeted by multiple pathogens during pathogenesis (Elmore and Coaker, 2011b). Previously, we reported that PM-associated plant CNLs utilized PMAs, including NbPMA3, by inhibiting their activity to promote cell death (Lee et al., 2022). However, whether pathogen effectors could promote cell death through modulation of PMA and the associated molecular mechanisms are still largely unknown.

To identify the functional RxLR effectors of *P. capsici*, a new pipeline was devised adopting three data sets, which included microarray expression profiles of a previous study (Jupe et al., 2013), open reading frame extraction (getORFs), and target domain-based annotation (TGFam-Finder) (Kim et al., 2020). A total of 268 and 217 putative RxLR effectors were isolated by getORFs and TGFam-Finder, respectively (Supplemental Figure S1). The candidates from these two datasets were merged with the public microarray expression data set, which lists RxLR genes upregulated during the biotrophic phase of *P. capsici* infection (Jupe et al., 2013). As a result, 25 putative RxLR effectors were selected, and each effector domain was artificially synthesized for further study (Supplemental Table S1).

Next, to screen for PMA-associated *P. capsici* RxLR effectors, the selected effectors from our multiple-omics screening were cloned into the potato virus X-based vector pICH31160 without epitope tag and used for *Agrobacterium*-mediated transient expression in *N. benthamiana* with or without 1 μM fusicoccin (FC), an irreversible PMA activator (Baunsgaard et al., 1998). Among them, 14 candidate effectors that induced cell death in *N. benthamiana* were named the <u>C</u>apsici <u>R</u>xLR cell death <u>I</u>nducer <u>S</u>uppressing <u>I</u>mmune <u>S</u>ystem (CRISIS). We observed that FC significantly inhibited the cell death induced by 8 effectors (CRISIS2, 3, 4, 5, 8, 10, 13, and 14) and enhanced the cell death induced by one effector (CRISIS 11) but had no effect on 5 other effectors (CRISIS1, 6, 7, 9, 12). The extent of cell death was photographed at 3 day-post-infiltration (dpi) and quantified as the quantum yield of photosystem II (PSII; Fv/Fm) (Figure 1A and 1B). To determine the association between NbPMA3 and FC-affected CRISIS, GFP protein was fused at the N-terminus of each effector, followed by a co-immunoprecipitation (co-IP) assay. Only CRISIS2 was identified as an interactor of NbPMA3 (Figure 1C). Subsequently, we focused on the characterization of CRISIS2, which exhibited compromised cell death by FC treatment and was physically associated with NbPMA3.

**Figure 1.**
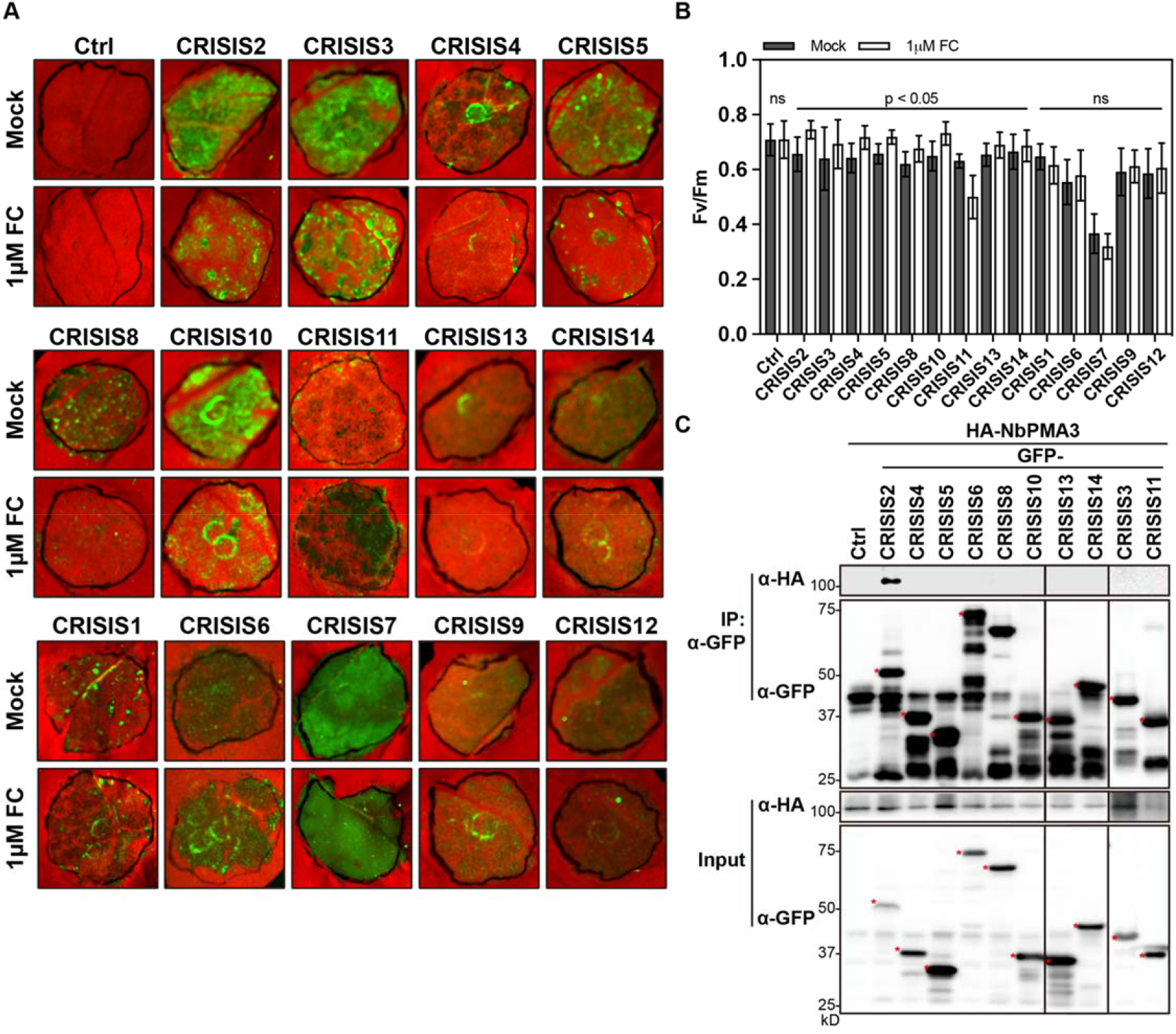
Functional screening of RxLR effector associated with PMA. **A**, Screening of fusicoccin effect on CRISIS-induced cell death. CRISISs were transiently overexpressed in *N. benthamiana*, followed by 1uM fusicoccin (FC) treatment at 16 hpi. The photographs were taken at 3 dpi. Empty vector (EV) was used as a negative control for cell death. **B**, The cell death intensity of A was quantified by measuring quantum yield (Fv/Fm). Data are represented as mean ± S.D (n ≥ 6). Significant difference between mock- or 1uM FC-treatment was determined using paired t-test. Ns, not significant. **C**, CRISIS2, but not other CRISISs interact with NbPMA3 *in planta*. HA-NbPMA3 was co-expressed with EV or GFP-CRISIS in *N. benthamiana* leaves. Proteins were extracted and subjected to immunoprecipitation (IP) with α-GFP agarose (IP: α-GFP) and immunoblotted with α-HA and α-GFP (top two panels). Input was collected from same protein extracts before IP (bottom two panels).

### CRISIS2-induced cell death is dose-dependent

During co-IP analyses, we observed that p35S:GFP-CRISIS2 lost cell death-inducing activity, we therefore reconstructed CRISIS2 with pCAMBIA2300 containing CaMV 35S promoter or potato virus X (PVX) based vector pKW (Lacomme and Chapman, 2008). FLAG or GFP were tagged at the N-termini of CRISIS2 in *p35S:FLAG-CRISIS2* (pCAMBIA2300) and *PVX:FLAG-CRISIS2* or *PVX:GFP-CRISIS2* (pKW) constructs (Figure 2A), given that C-terminal tag fusion is known to interfere with the function of effector (Bos et al., 2006). The cell death-inducing activity was observed in *N. benthamiana* after infiltration with *Agrobacterium* having each construct. Interestingly only PVX:FLAG-CRISIS2 was able to induce cell death (Figure 2B, top panel). The expression of CRISIS2 was potentiated at 2 dpi in the leaves infiltrated with PVX:FLAG-CRISIS2 while p35S:FLAG-CRISIS2 was steadily expressed through 1 to 3 dpi. GFP tagged constructs in both vector system were weakly expressed (Figure 2B, bottom panel). We further confirmed that only *PVX:FLAG-CRISIS2* promoted the expression of cell death marker genes, *NbHIN1*, *NbHsr203J*, and *NbWIPK* (Figure 2C) (Melech-Bonfil and Sessa, 2010; Moon et al., 2016). These results indicate that the cell death induced by CRISIS2 is dose-dependent. CRISIS2 protein accumulation over the certain threshold is therefore likely required to induce cell death.

**Figure 2.**
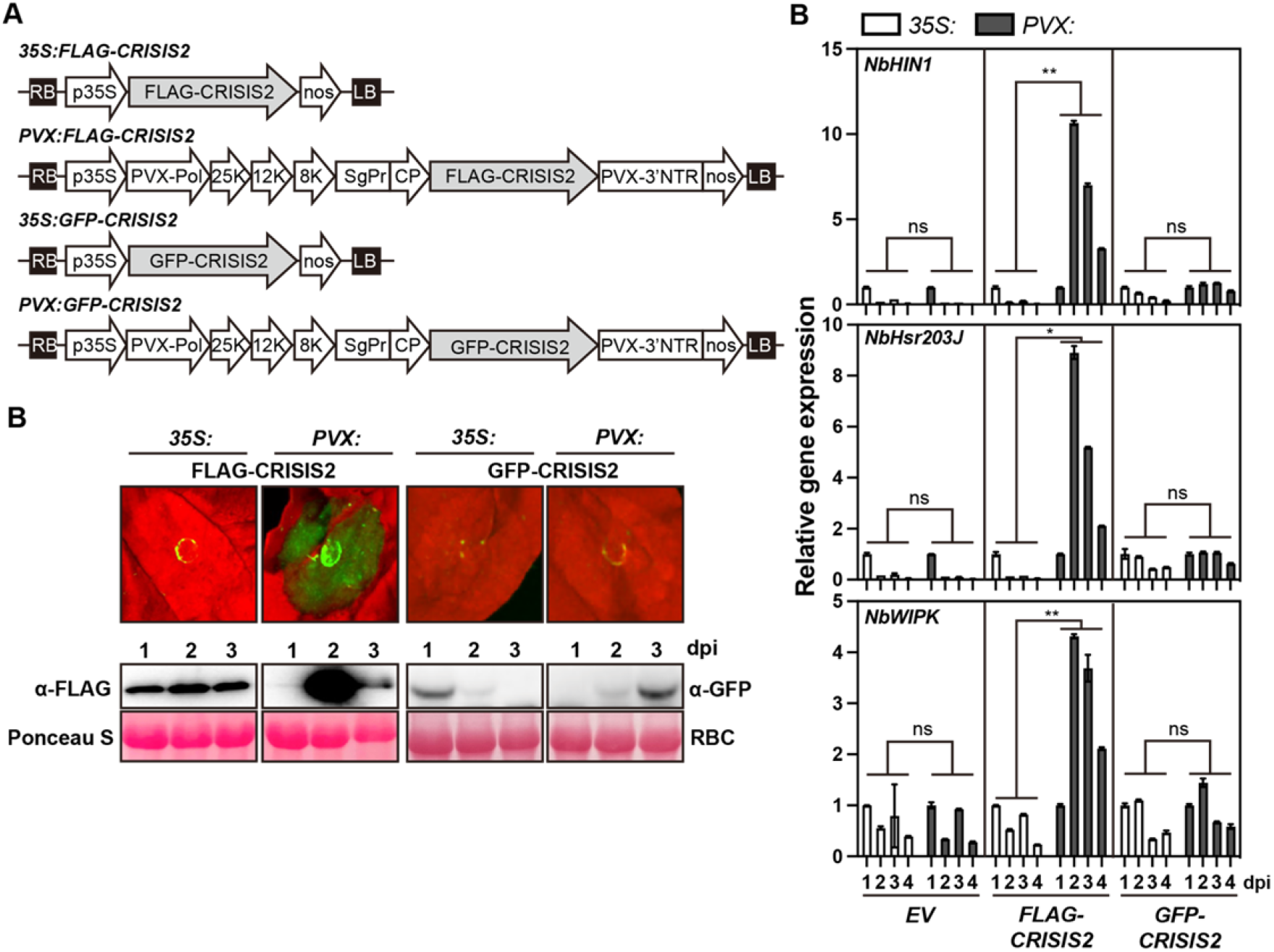
CRISIS2-induced cell death is dose-dependent. **A**, Schematic representations of the T-DNA constructs. LB, left border; RB, right border; p35S, CaMV 35S; nos, *A. tumefaciens* nopaline synthetase gene terminator; PVX-Pol, polymerase from Potato virus X (PVX); 25K, 12K, 8K: PVX triple gene block; SgPr: subgenomic promoter; CP: PVX coat protein; PVX-3′-NTR: PVX 3′ nontranslated region. **B**, CRISIS2-induced cell death is dose-dependent. FLAG-CRISIS2 or GFP-CRISIS2 was transiently expressed in *N. benthamiana* leaves under the control of 35S promoter (35S:FLAG-CRISIS2 or 35S:GFP-CRISIS2) or PVX (PVX:FLAG-CRISIS2 or PVX:GFP-CRISIS2). The photograph was taken at 3 dpi (top panel). The protein accumulation at 1, 2, and 3 dpi was detected by immunoblotting with α-FLAG or α-GFP (middle panel). Ponceau S staining of rubisco (RBC) is shown as the loading control (bottom panel). **C**, Relative expression levels of cell death marker genes of *N. benthamiana* upon transient expression of *35S:EV, 35S:FLAG-CRISIS2, 35S:GFP-CRISIS2, PVX:EV, PVX:FLAG-CRISIS2*, or *PVX:GFP-CRISIS2* by agro-infiltration. The tissues were harvested at each time point, followed by RNA extraction. Transcript levels of each gene were determined by qRT-PCR, which is normalized to *NbEF1⍺* gene. Data are normalized to 1 dpi of each construct and presented as mean ± SD (n ≥ 2). Significant difference was analyzed by one-way ANOVA (Tukey’s multiple comparisons test). ns, not significant; *, p < 0.05; **, p < 0.01.

### CRISIS2 interacts with NbPMAs at the plasma membrane and CRISIS2-induced cell death is inhibited by NbPMAs

To further demonstrate the FC effect on CRISIS2-induced cell death, we expressed the PVX:FLAG-CRISIS2 in *N. benthamiana* leaves using agroinfiltration followed by FC treatment at 12 hour post infiltration (hpi). As expected, the cell death induced by CRISIS2 was significantly reduced in FC-treated leaves without any significant change of protein expression (Figure 3A-C). PVX:FLAG-CRISIS6, which has cell death-inducing activity but is not affected by FC treatment, was used as a negative control. Next, the subcellular localization of CRISIS2 and CRISIS6 was determined by confocal microscopy. The fluorescence signal of GFP-CRISIS2 was solely detected at the PM in control and plasmolyzed cells (Figure 3D upper two panels), while multiple signals from the PM and cytosol were detected in both control and plasmolyzed cells expressing GFP-CRISIS6 (Figure 3D lower two panels). PM localization of CRISIS2 was additionally confirmed by co-localization with NbPMA3-mStrawberry (Supplemental Figure S2). Moreover, FC did not affect the localization of either CRISIS2 or CRISIS6 (Figure 3E right two panels). These results indicated that FC-induced inhibition of cell death was not due to defective expression or mis-localization of CRISIS2. Together, our observations demonstrate that CRISIS2 is a novel and biologically functional RxLR effector with cell death-inducing activity at the PM of the host cell.

**Figure 3.**
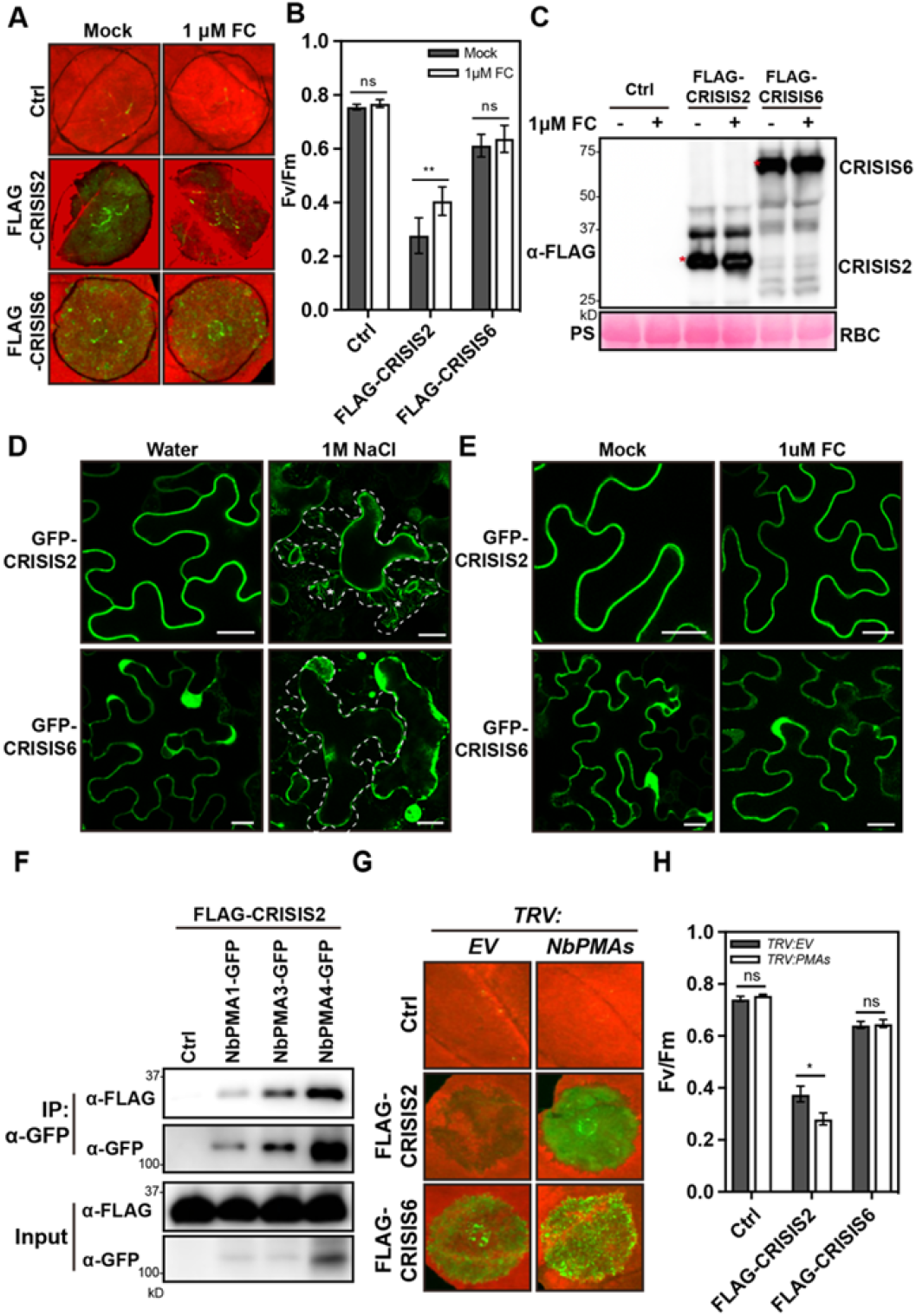
CRISIS2 interacts with NbPMAs at PM and CRISIS2-induced cell death is inhibited by NbPMAs. **A**, The PMA activator fusicoccin (FC) attenuates cell death induced by CRISIS2. Control empty vector (Ctrl), FLAG-CRISIS2 and FLAG-CRISIS6 were transiently expressed in *N. benthamiana* leaves by agro-infiltration. At 16 hours post-infiltration (hpi), 1 μM FC was infiltrated into the agro-infiltrated region. The leaves were photographed at 3 days post infiltration (dpi). **B**, The cell death intensity in (A) was quantified by measuring quantum yield (Fv/Fm). Data are represented as mean ± SD (n = 6). Significant differences were determined using unpaired t-tests. ns, not significant; **, p-value < 0.01. **C**, Protein accumulation of CRISIS2 and CRISIS6 in mock- or 1 μM FC-treated *N. benthamiana*. Proteins were extracted and subjected to immunoblotting with α-FLAG. Ponceau S staining of rubisco (RBC) is shown as the loading control. **D**, CRISIS2 localizes to plasma membrane. Cells expressing GFP-CRISIS2 and GFP-CRISIS6 were plasmolyzed with 1 M NaCl solution. The dashed lines indicate the cell walls. Asterisks indicate Hechtian strands. Bar = 20 μm. **E**, Fusicoccin does not affect the subcellular localization of GFP-CRISIS2. Agro-infiltrated leaves were treated with 1 μM FC at 16 hpi. Scale bar = 20 μm. Confocal microscopy images were taken at 48 hpi. **F**, CRISIS2 was co-expressed with EV (Ctrl), NbPMA1-GFP, NbPMA3-GFP, or NbPMA4-GFP in *N. benthamiana* leaves. Proteins were extracted and subjected to immunoprecipitation (IP) with α-GFP agarose (IP: α-GFP) and immunoblotted with α-FLAG and α-GFP (top two panels). Input was collected from same protein extracts before IP (bottom two panels). **G**, Cell death induced by CRISIS2 is enhanced by *NbPMAs* silencing. *Agrobacteria*-carrying EV (Ctrl), FLAG-CRISIS2, or FLAG-CRISIS6 were infiltrated into *EV*- or *NbPMAs-*silenced *N. benthamiana* leaves. Photographs were taken at 3 dpi. **H**, The cell death intensity in (G) quantified by measuring quantum yield (Fv/Fm). Data are represented as mean ± SE (n = 33). Significant difference was analyzed by unpaired t-test. ns, not significant; *, p-value < 0.05.

Initially, we hypothesized that if the cell death-inducing activity of RxLR effectors is affected by FC, these effectors would be localized at the PM. However, only CRISIS2, CRISIS3, and CRISIS8 among the 10 putative CRISISs co-localized with NbPMA3 (Supplemental Figure S2). The localization of the other candidates only partially overlapped with NbPMA3, as additional GFP fluorescence signals were detected from the cytosol and/or nucleus. These results suggest that FC has likely unknown indirect effects besides affecting RxLR-mediated cell death through PM-localized PMAs.

Previously, we reported a functional redundancy of several NbPMAs in *N. benthamiana* (Lee et al., 2022). Co-IP revealed that CRISIS2 also associated with NbPMA1 and NbPMA4, implying that CRISIS2 functions with multiple NbPMAs (Figure 3F). To further demonstrate the functional association between CRISIS2 and NbPMAs, first, CRISIS2 or CRISIS6 were co-expressed with NbPMA3 in *N. benthamiana* to observe and quantify cell death. CRISIS2-induced cell death was slightly inhibited by transient overexpression of NbPMA3 but not significantly (Supplemental Figure S3). However, remarkably enhanced cell death and decreased quantum yield by CRISIS2, but not by CRISIS6, were observed in either NbPMA3 (data not shown) or *NbPMA1/3/4*-silenced plants using virus-induced gene silencing (VIGS) compared with the control plants (Figure 3G and 3H). The protein expression and gene silencing efficiency were confirmed by immunoblot analysis and quantitative reverse transcription PCR (qRT-PCR), respectively (Supplemental Figure S4A and S4B). These results suggest that NbPMAs function as negative regulators of CRISIS2-induced cell death.

### CRISIS2-induced cell death is independent on NbSGT1 and NbRAR1

NLR proteins are major intracellular receptors recognizing pathogen effectors and subsequently induce HR cell death to quarantine the site infected by pathogen. To determine whether CRISIS2-induced cell death is dependent on NLR, we silenced *NbSGT1*, *NbRAR1*, or *NbHSP90*, the core regulatory components of NLR-mediated HR cell death, in *N. benthamiana* using VIGS (Azevedo et al., 2002,(Botër et al., 2007). CRISIS2, CRISIS6, Rpiblb2-Avrblb2 (Potato R genes and *Phytophthora infestans* RxLR effector, respectively) as a positive control, necrosis-inducing protein (NIP) as a negative control (Oh et al., 2014), or EV were transiently expressed in *EV*- or *NbSGT1*-silenced plants. As expected, we observed that the cell death induced by Rpiblb2-Avrblb2 was compromised in *NbSGT1-*silenced plants, whereas NIP-induced cell death was not affected by *NbSGT1* silencing. In contrast, similar extent of cell death and Fv/Fm were observed in CRISIS2-expressing leaves in EV- or *NbSGT1*-silenced plant (Figure 4A and 4B). Moreover, the silencing of *NbRAR1* also did not affect CRISIS2-induced cell death (Figure 4E and 4F). CRISIS6-induced cell death was compromised in *NbSGT1*-silenced plant but was not affected in *NbRAR1*-silenced plant, as observed for Rpiblb2-Avrblb2-induced cell death (Oh *et al*., 2014). We failed to obtain significant data in *HSP90*-silenced plant due to severe developmental defect (data not shown). The expression of CRISIS2 or CRISIS6 proteins and the silencing efficiency of *NbSGT1* or *NbRAR1* were determined immunoblot (Figure 4C and 4G) or qRT-PCR (Figure 4D and 4H), respectively. Together, these results imply that CRISIS2*-*induced cell death is not mediated by NLR(s) which requires SGT1 or RAR1, suggesting that CRISIS2 triggers cell death through a distinct mechanism.

**Figure 4.**
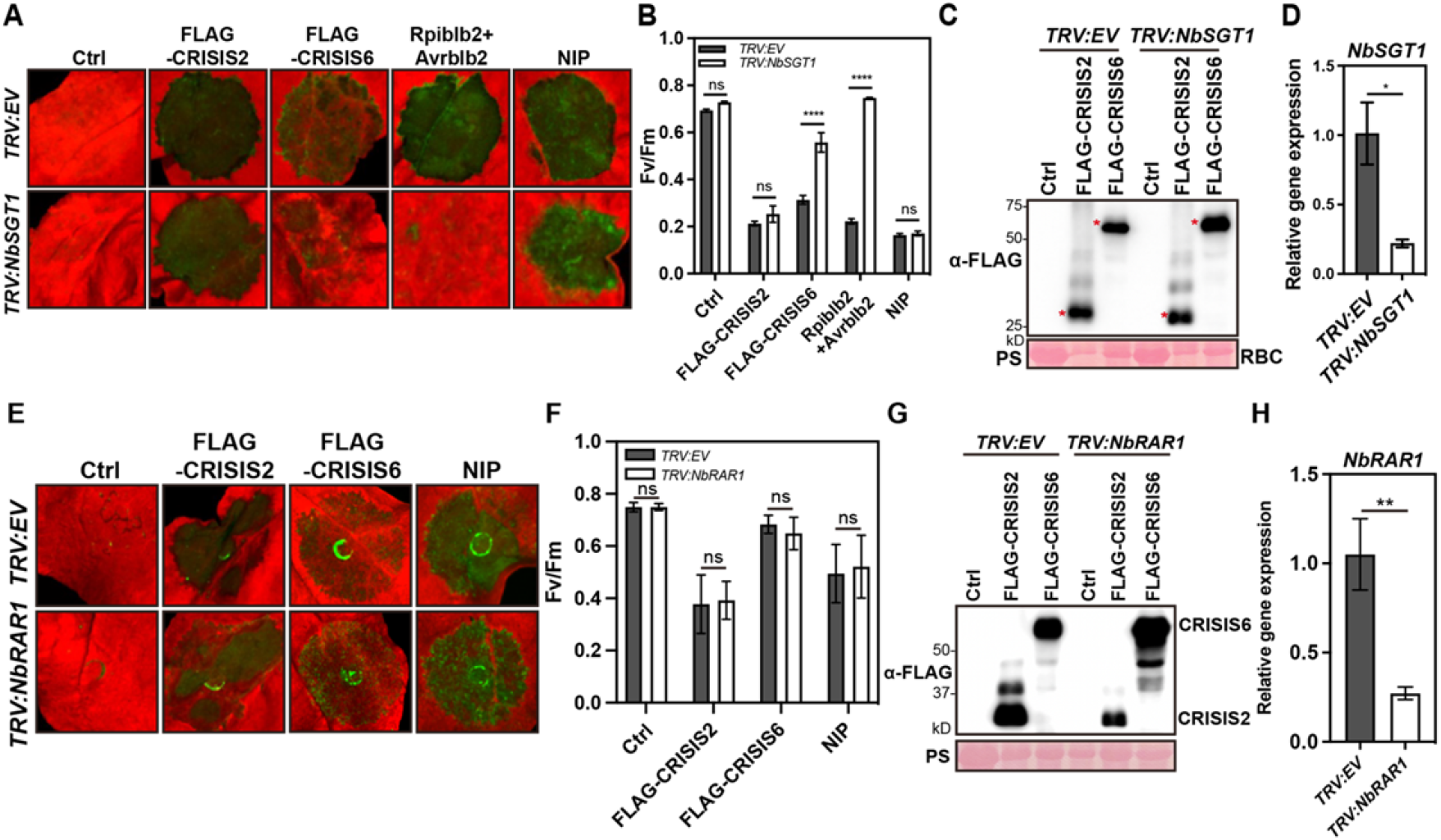
CRISIS2-induced cell death is independent on NbSGT1 and NbRAR1. **A**, Cell death induced by CRISIS2 is independent of *NbSGT1*. Agrobacteria carrying EV (Ctrl), FLAG-CRISIS2, FLAG-CRISIS6, Rpiblb2+Avrblb2, or NIP were infiltrated in *EV*- or *NbSGT1*-silenced *N. benthamiana*. Photographs were taken at 3 dpi. Rpiblb2+Avrblb2 and NIP were used as *NbSGT1*-dependent and *NbSGT1*-independent cell death controls, respectively. **B**, The cell death intensity in (A) quantified by measuring quantum yield (Fv/Fm). Data are represented as mean ± SE (n ≥ 15). Significant difference was analyzed by one-way ANOVA (Tukey’s multiple comparisons test). ns, not significant; ****, p < 0.0001. **C**, Accumulation of of FLAG-CRISIS2 and FLAG-CRISIS6 proteins expressed in *TRV:EV* and *TRV:NbSGT1* plants in (G). Ponceau S (PS) staining of rubisco (RBC) is shown as the loading control. **D**, Transcript accumulation measured by qRT-PCR to confirm the silencing efficiency of *NbSGT1*. The transcript accumulation of *NbSGT1* was measured at 2 weeks after VIGS. Data are represented as mean ± SD. Significant differences were determined by unpaired t-tests. *, p < 0.05. **E**, Cell death induced by CRISIS2 is independent of *NbRAR1*. Agrobacteria carrying EV, FLAG-CRISIS2, FLAG-CRISIS6, or NIP were infiltrated in *EV*- or *NbRAR1*-silenced *N. benthamiana*. Photographs were taken at 3 dpi. NIP was used as *NbRAR1*-independent cell death control. **F**, Cell death intensity in (E) quantified by measuring quantum yield (Fv/Fm). Data are represented as mean ± SD (n ≥ 9). Significant difference was determined using one-way ANOVA. ns, not significant. **G**, Accumulation of FLAG-CRISIS2 and FLAG-CRISIS6 proteins expressed in *TRV:EV* and *TRV:NbRAR1* plants in (E). Ponceau S staining of rubisco (RBC) is shown as the loading control. **H**, Transcript accumulation measured by qRT-PCR to confirm the silencing efficiency of *NbRAR1*. The transcript accumulation of *NbRAR1* was measured at 2 weeks after VIGS. Data are represented as mean ± SD. Significant difference was determined by unpaired t-test. **, p-value < 0.01.

### CRISIS2 induces alkalization of apoplasts by inhibiting PMA activity

Apoplastic alkalization due to modification of PMA activity often leads to cell death (Chen et al., 2010; Fuglsang, 2020). To monitor the change of pH in the apoplast, *A. tumefaciens* carrying *FLAG-CRISIS2* or EV were infiltrated into ratiometric pHluorin sensor (PM-APO)-transgenic *N. benthamiana* leaves (Martiniere et al., 2018). The fluorescence ratio was monitored in the epidermal cells, and the corresponding pH values were calculated using *in vitro* calibration with a recombinant pHluorin (Supplemental Figure S5). The apoplastic pH dramatically increased from 1.5 to 2 dpi in *FLAG-CRISIS2*-infiltrated leaves but not in the EV control leaves (Figure 5A and 5B). Considering that CRISIS2-induced cell death appears after 2.5 dpi, this result suggests that PMA activity is inhibited by the expression of CRISIS2 before the onset of macroscopic cell death. To determine whether CRISIS2 indeed inhibits PMA activity, we measured PMA activity *in vivo*. Due to technical limitations in measuring the biochemical activity of single PMA isoforms in the plant, we analyzed PMA activity in whole leaf extracts of CRISIS2-expressing or EV control *N. benthamiana*. The leaf samples were harvested at 2 dpi, and PM vesicles were prepared for measurement of ATPase activity (Palmgren, 1990). The ATPase activity in *FLAG-CRISIS2*-expressing leaves was significantly lower than that in the EV control leaves (Figure 3C). The efficiency of membrane fractionation was confirmed by western blot analyses of fractionated samples with specific antibodies for PM-localized PMA and the cytosolic phosphoenol pyruvate carboxylase (PEPC) (Supplemental Figure S6A). Because the ATPase activity was measured by detecting inorganic phosphate in solution, we examined whether CRISIS2 associates with other P-type ATPases, such as the calcium-transporting ATPase NbACA8, localized in PM (Yu et al., 2018). However, CRISIS2 interacted only with NbPMA3, not with NbACA8 (Supplemental Figure S6B). These results indicate that the decreased ATPase activity likely stemmed from the interaction between CRISIS2 and H^+^-ATPase but not Ca^2+^-ATPase at PM. We further confirmed that the ATPase activity was dramatically compromised by the cell death-inducing PVX:FLAG-CRISIS2 but only marginally affected by the p35S:FLAG-CRISIS2 construct that lacks the cell death-inducing activity (Supplemental Figure S6C). This result indicates that CRISIS2-induced cell death is linked with the inhibition of PMA activity.

**Figure 5.**
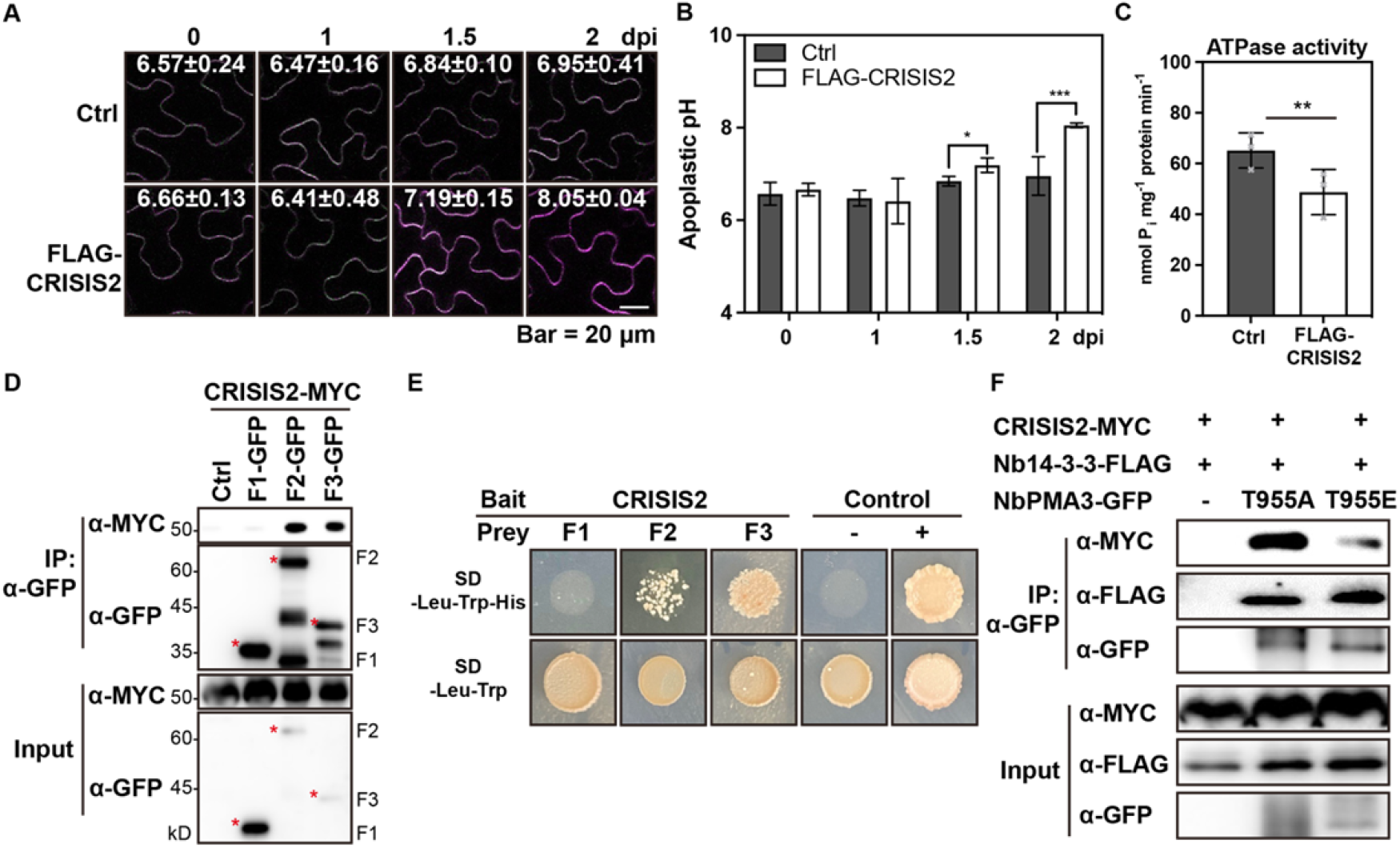
CRISIS2 induces apoplastic alkalization by inhibiting PMA. **A**, CRISIS2-induced apoplastic alkalization. The apoplastic pH of *EV* (*Ctrl*)- or *FLAG-CRISIS2*-expressing *N. benthamiana* leaf epidermal cells was monitored using the apoplastic pH sensor PM-Apo. The mean ± SD of the calculated pH is shown in each image. Images were taken under the same microscope settings at 0, 1, 1.5, and 2 dpi. The pH was calculated using an *in vitro* calibration with recombinant pHluorin (see Supplemental Figure 6). Bar = 20 μm. **B**, The pH values of (A). Data are represented as mean ± SD (n ≥ 4). Significant differences were determined by two-way ANOVA. *, p-value < 0.05; ***, p-value < 0.001. **C**, Decreased ATPase activity in FLAG-CRISIS2-expressing *N. benthamiana* leaves. Microsomal fractions isolated from *N. benthamiana* leaves expressing EV or FLAG-CRISIS2 were used to measure PMA activity. Data are represented as mean ± SD (n = 3). Significant difference was determined by paired t-test. The experiment was performed three times with similar results. **D**, CRISIS2 interacts with C-terminal of NbPMA3. CRISIS2-MYC was co-expressed into *N. benthamiana* leaves with EV or three cytosolic domains of NbPMA3 (F1, 1 – 64 residues; F2, 305 – 650 residues; F3, 846 – 956 residues; see Supplemental Figure S7). Extracted protein was subjected to immunoprecipitation (IP) with α-GFP agarose (IP: α-GFP) and immunoblotted with α-MYC and α-GFP (top two panels). Input was collected from the same protein extracts before IP (bottom two panels). **E**, CRISIS2 interacts with the F2 and F3 cytosolic domains of NbPMA3 in yeast. CRISIS2 was cloned into the bait plasmid pGBKT7, and the F1, F2, and F3 domains of NbPMA3 were cloned into the prey plasmid pGADT7. A combination of Nb14-3-3 (Bait) and the F3 domain of NbPMA3 (Prey) was used as a positive control, and combinations of Lam (Bait) and the F1, F2, and F3 domains of NbPMA3 (Prey) were used as negative controls. The presented image is a representative data. Yeast transformants were grown on SD/-Leu/-Trp and selected on SD/-Leu/-Trp/-His. The plates were photographed 7 days after plating. **F**, CRISIS2 associates with phosphor-null NbPMA3 (T955A) more strongly than with phosphor-mimic NbPMA3 (T955E) *in planta*. CRISIS2-MYC and Nb14-3-3-FLAG were co-expressed with EV, GFP-NbPMA3 (T955A) or GFP-NbPMA3 (T955E) in *N. benthamiana* leaves. Proteins were extracted and subjected to immunoprecipitation (IP) with α-GFP agarose (IP: α-GFP) and immunoblotted with α-MYC, α-FLAG and α-GFP (top three panels). Input was collected from the same protein extracts before IP (bottom three panels). The experiments were performed two times with similar results.

Since CRISIS2 is a cytoplasmic effector associated with PM-localized PMAs, we investigated how CRISIS2 could regulate PMA activity. PMAs contain three major cytosolic domains, including the N-terminal actuator domain (here fragment F1, Supplemental Figure S7), a central catalytic domain (F2), and a C-terminal autoinhibitory domain (F3). To determine which domain(s) of PMA could associate with CRISIS2, co-IP and yeast two-hybrid assays were performed with the individual NbPMA3 cytosolic domains. CRISIS2 was strongly associated with the F2 and F3 domains in both assays (Figure 5D and 5E). The F3 domain appeared to be the major binding domain of CRISIS2 based on the growth of yeast. These results suggest that CRISIS2 possibly regulates the activation status of PMAs by interacting with the C-terminal regulatory domain of PMAs.

The activation of PMAs is mainly regulated by posttranslational modifications, such as the phosphorylation of various serine (Ser) and threonine (Thr) residues in the C-terminal regulatory domain (Haruta et al., 2015). Once PMA is activated by phosphorylation at its penultimate Thr residue, it allows for the association of the activator protein 14-3-3 and confers the transition from the autoinhibition to activation state (Elmore and Coaker, 2011; Jelich-Ottmann et al., 2001). To test the association between CRISIS2 and NbPMA3 presenting altered activation status, phosphomimic (T955E) and null-phospho (T955A) mutants at the penultimate Thr residue of NbPMA3 were constructed, and a co-IP assay was performed with CRISIS2. Nb14-3-3 was added to confirm the phospho state of the NbPMA3 variants. As expected, the 14-3-3 protein exhibited a greater association with the phosphomimic (T955E) mutant than with the null-phospho (T955A) mutant (Figure 3F upper middle panel). However, a significantly stronger interaction was observed between CRISIS2 and the null-phospho (T955A) variant (Figure 3F upper top panel). This was reminiscent of the PM-associated plant CNLs that inhibit PMAs activity to promote cell death (Lee et al., 2022). This result suggests that CRISIS2 associates with inactive PMA, potentially preventing PMA activation. Altogether, our data demonstrates that CRISIS2 can inhibit the activity of PMAs by binding to the dephosphorylated penultimate Thr residue, hence probably preventing the transition of inactive to activated PMA.

### CRISIS2 positively affects to the virulence of *P. capsici* in *N. benthamiana*

As a next step, we determined the contribution of CRISIS2 to the virulence of *P. capsici*. First, we measured the expression pattern of CRISIS2 during *P. capsici* infection in *N. benthamiana* using qRT-PCR and found that the transcripts of both CRISIS2 and CRISIS6 were dramatically induced at 6 h post inoculation (hpi), which is similar to the expression pattern of the biotrophic phase marker gene *PcHmp1* (*P. capsici* haustorial membrane protein1) (Avrova et al., 2008) (Figure 6A). The expression of the necrotrophic phase marker gene, *PcNPP1* (*P. capsici necrosis-inducing Phytophthora protein 1*), was evident at 18-48 hpi during the transition from biotrophic to necrotrophic stage following pathogen infection (Jupe et al., 2013). To determine the role of CRISIS2 in pathogenicity, while considering the undistinguishable symptoms from CRISIS-induced cell death and pathogen-induced necrotic lesion, we expressed p35S:*GFP-CRISIS2, p35S:GFP-CRISIS6* (both defective for cell death-inducing activity) or p35S:*GFP* in *N. benthamiana* 24 h before *P. capsici* inoculation. Significantly enlarged disease lesions were observed in CRISIS2*-* but not in CRISIS6-expressing leaves compared to the GFP control (Figure 6B). Protein expression was confirmed by western blot (Supplemental Figure S8A).

**Figure 6.**
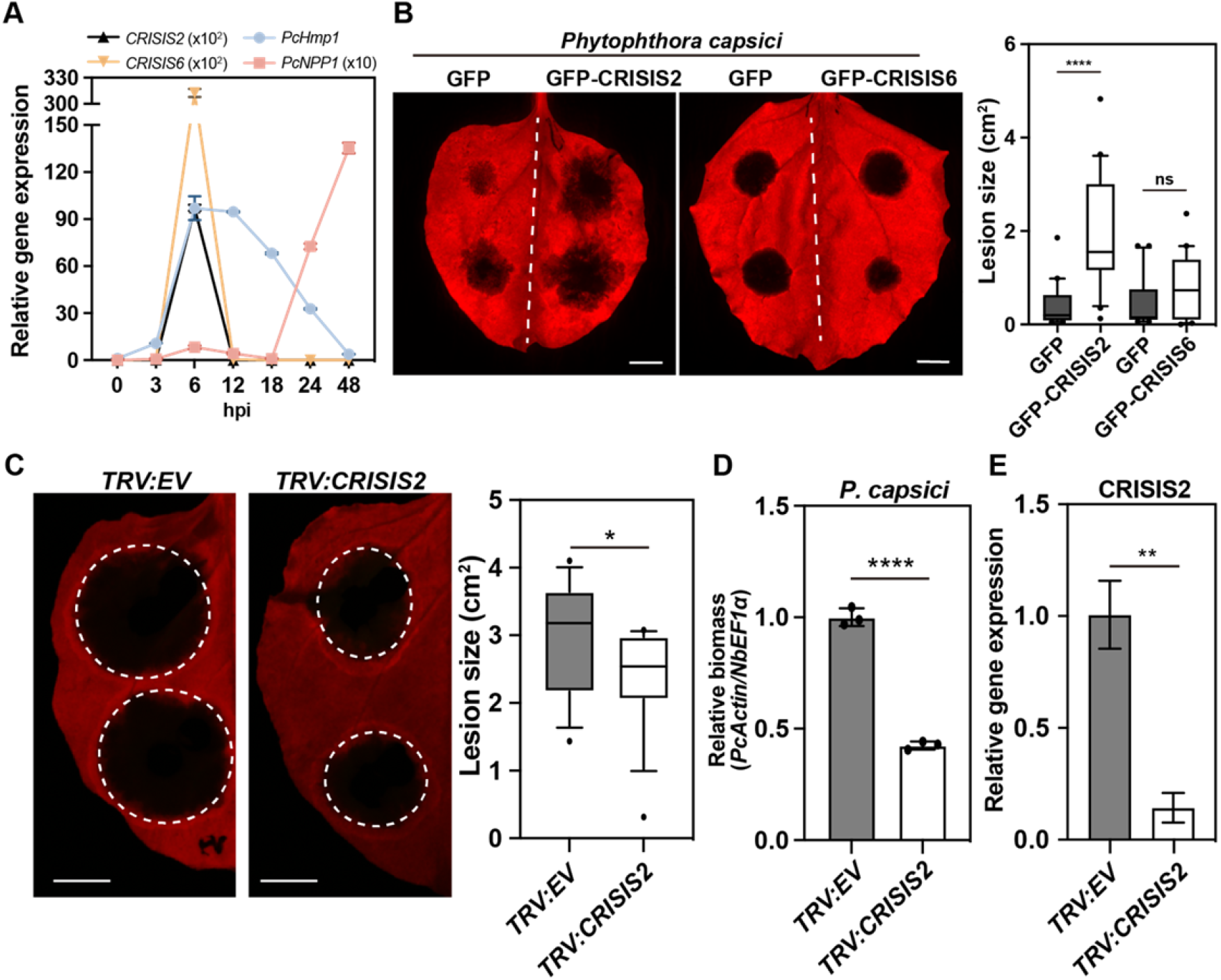
CRISIS2 contributes to the virulence of *Phytophthora capsici*. **A**, Expression profiles of *CRISIS2*, *CRISIS6*, *PcHmp1*, and *PcNPP1*. Leaves of *N. benthamiana* were inoculated with *P. capsici*, and the infected tissue were harvested at each time point, followed by RNA extraction. Transcript levels of *CRISIS2*, *CRISIS6*, *PcHmp1*, and *PcNPP1* were measured by qRT-PCR, which is normalized to *PcTubulin* gene. **B**, GFP, GFP-CRISIS2 or GFP-CRISIS6 were expressed in *N. benthamiana* leaves by agro-infiltration, followed by *P. capsici* inoculation at 24 hpi. The leaves were photographed 2 days after *P. capsici* inoculation (left panel). The lesion areas were measured and quantified (right panel). The data are represented as 10-90% boxes and whiskers (n = 20). Significant differences were determined by unpaired t-tests. ns, not significant; ****, p-value < 0.001. **C**, Host-induced *EV-* or *CRISIS2-*silenced *P. capsici* inoculation on plants (left panel). *N. benthamiana* were treated with TRV vectors harboring *EV* or *CRISIS2* gene fragment by agro-infiltration. After 10 days of TRV treatment, *P. capsici* was drop-inoculated in the upper leaves. The photographs were taken at 2 days after *P. capsici* infection. The lesion areas were measured and quantified (right panel). The data are represented as 10-90% boxes and whiskers (n = 16). Significant difference was determined by unpaired t-tests. *, p-value < 0.05. **D**, Biomass of *P. capsici* in (C). At 2 days after *P. capsici* inoculation, leaf disks around the inoculated site were harvested. The total genomic DNA was extracted and subjected to qPCR analysis. The biomass of *P. capsici* was determined by measuring *PcActin* gene normalized to *NbEF1⍺* gene. The significant difference compared with *TRV:EV* was determined by unpaired t-test. ****, p < 0.0001. **E**, Relative transcript levels of *CRISIS2* in host-induced *EV-* (*TRV:EV*) or *CRISIS2-*silenced (*TRV:CRISIS2*) *P. capsici*. After 6 h *P. capsici* inoculation, the inoculated regions of leaves were harvested and subjected to qRT-PCR analysis. The transcript level of *CRISIS2* was normalized to *PcTubulin* gene. Significant difference was determined by unpaired t-tests. **, p-value < 0.01.

Alternatively, we used host-induced gene silencing (HIGS) to impair CRISIS2 expression in *P. capsici* (Zhu et al., 2017) (Figure 6C and 6D). Consistently with CRISIS2-overexpression study, we also observed smaller necrotic lesions and reduced pathogen biomass, an indication of pathogen growth, in leaves infected with CRISIS2-silenced *P. capsici*. Silencing efficiency of CRISIS2 was confirmed by measuring the relative transcript level of *CRISIS2* at 6 h after *P. capsici* infection (Figure 6E). These results indicate that CRISIS2 is indeed required for the full virulence of *P. capsici* in host plants.

### CRISIS2 suppresses PTI responses

Adapted pathogens have acquired effector proteins that are secreted into the plant cell where they suppress PTI (Block et al., 2008). Since rapid MAPK activation and production of ROS are critical signaling components triggering PTI (Kadota et al., 2014; Keinath et al., 2010), flg22-induced MAPK activation and ROS production were measured in *N. benthamiana* following expression of CRISIS2 or CRISIS6. In the EV expressing leaves, NbMAPKs, SIPK and WIPK (Yang et al., 2001) were activated by flg22 at 15 min after treatment and the induction was reduced at 45 min after treatment (Figure 7A and 7B top panel). However significantly reduced activation of NbSIPK and NbWIPK was observed in CRISIS2-expressing leaves at 15 and 30 min after treatment, while no difference was observed in CRISIS6-expressing leaves. CRISIS2 and CRISIS6 expression was confirmed by western blot (Figure 7A and 7B middle panel). Similarly, significant reduction of ROS production in response to flg22 was observed in CRISIS2-expressing leaves compared to the CRISIS6- or GFP-expressing control (Figure 7C). Protein accumulation of CRISIS2 and CRISIS6 was confirmed by western blot analysis of the leaf samples used for the ROS assay (Supplemental Figure S8B). Consistent with these data, significantly reduced expression of the early PTI marker gene *NbCYP1D20* (Heese et al., 2007) and late-expressed defense-related genes *NbPR1* and *NbWRKY8* was observed in CRISIS2-expressing leaves of *N. benthamiana* upon *P. capsici* infection (Figure 7D). Moreover, a lower ROS production and reduced defense-related gene transcript accumulation were also observed in CRISIS2-expressing tissues in response to the unrelated PAMP elf18, an 18-amino acid N-terminal epitope of the bacterial elongation factor Tu recognized by EFR (Kunze et al., 2004) (Supplemental Figure S9A-9C). Taken together, these data indicate that the RxLR effector CRISIS2 impairs PTI signaling, upstream of MAPK activation and ROS production.

**Figure 7.**
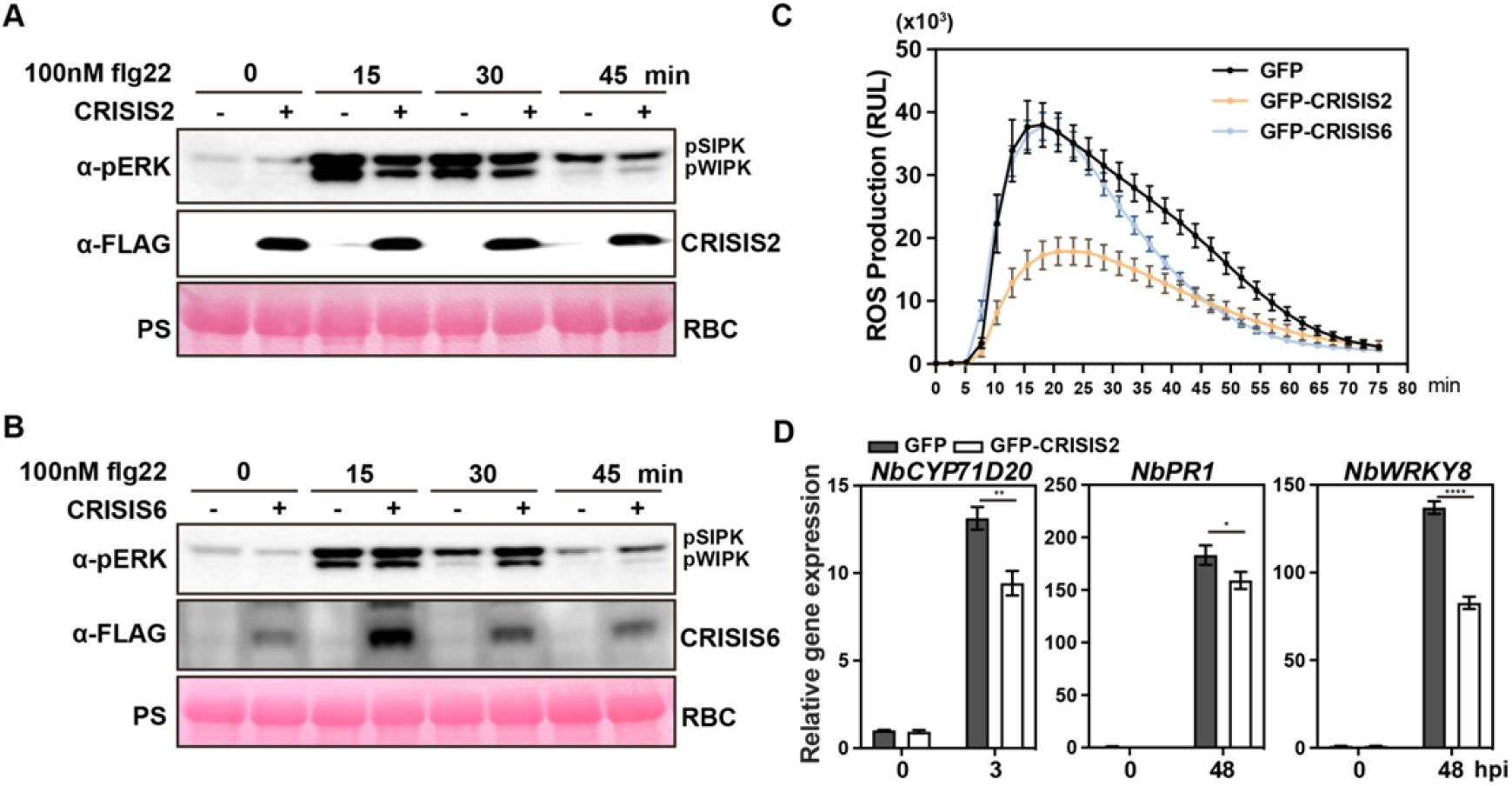
CRISIS2 inhibits PTI responses. **A-B**, CRISIS2 suppresses MAPK activation triggered by flg22. EV, FLAG-CRISIS2 (A), or FLAG-CRISIS6 (B) was expressed in *N. benthamiana* leaves. At 36 hpi, 100 nM flg22 was added for 0, 15, 30, and 45 min. Proteins were extracted, and MAPK activation was examined by α-pERK immunoblotting. Protein accumulation of FLAG-CRISIS2 and FLAG-CRISIS6 was detected by α-FLAG immunoblot. Ponceau S (PS) staining of rubisco (RBC) is shown as the loading control. **C**, CRISIS2 inhibits flg22-induced oxidative burst. GFP, GFP-CRISIS2 and GFP-CRISIS6 were transiently expressed in *N. benthamiana* leaves by agro-infiltration. At 2 dpi, ROS production was examined following treatment with flg22. Data are presented as mean ± SE from 24 leaf discs. **D**, Transient expression of CRISIS2 significantly reduces the expression of defense-related genes induced by *P. capsici* infection in *N. benthamiana*. GFP or GFP-CRISIS2 was expressed in *N. benthamiana* leaves by agro-infiltration, and then *P. capsici* was inoculated after 24 h. The transcript levels of *NbCYP1D20*, *NbPR1*, and *NbWRKY8* were measured by qRT-PCR at 3 h or 48 h after *P. capsici* inoculation. Data are normalized to 0 hpi and presented as the mean ± SD. Significant differences were determined by two-way ANOVA (Sidak’s multiple comparisons test). *, p < 0.05; **, p < 0.01; ****, p < 0.0001.

### CRISIS2 inhibits the flg22-induced FLS2-BAK1 association

Since CRISIS2 is localized at the PM and impairs flg22-induced ROS production and MAPK activation, it appears to function at a very early stage in PTI signaling. To test this hypothesis, co-IP assays were performed with CRISIS2 and the components of a pattern recognition receptor complex. Co-IP assays revealed that CRISIS2 associated with the PM-localized flg22 receptor FLS2 complex, including the co-receptor BAK1 (Figure 8A and 8B), but not with other PM proteins, such as NbCNGC4 and NbAUX1 (Supplemental Figure S10A), indicating that the interaction between CRISIS2 and the NbFLS2-NbBAK1 complex is specific. Perception of flg22 triggers the rapid formation of a complex of FLS2 and BAK1 (Chinchilla et al., 2007; Heese et al., 2007). In our assay, flg22 treatment markedly induced the interaction between NbFLS2 and NbBAK1, but the interactions between CRISIS2 and FLS2 or CRISIS2 and BAK1 were not affected (Figure 8B). However, the co-expression of CRISIS2 antagonized the flg22-induced FLS2-BAK1 association (Figure 8C). Next, we performed a virulence test in *NbBAK1*-silenced *N. benthamiana*. As expected, significantly enhanced susceptibility to *P. capsici* was observed in *NbBAK1*-silenced plants compared with the EV control (Chaparro-Garcia et al., 2011) (Figure 7D-7F). The enhanced virulence of *P. capsici* and decreased expression of *AtFRK1* (*FLG22-INDUCED RECEPTOR-LIKE KINASE 1*) were further confirmed in Arabidopsis *fls2* mutant compared with wild type Col-0 (Supplemental Figure S10B-S10D). However, we could not detect any difference of CRISIS2-induced cell death in *NbFLS2-*silenced *N. benthamiana* compared to *EV-*silenced plants, further supporting our hypothesis that CRISIS2 triggers cell death through PMA inhibition (Supplemental Figure S10E-10G). Taken together, these results indicate that CRISIS2 is closely associated with FLS2 and BAK1 in the resting state and inhibits ligand-induced FLS2-BAK1 receptor complex formation in response to flg22, thus negatively regulates early PTI signaling. This evidence also supports the contribution of CRISIS2 to the virulence of *P. capsici* in its host plant.

**Figure 8.**
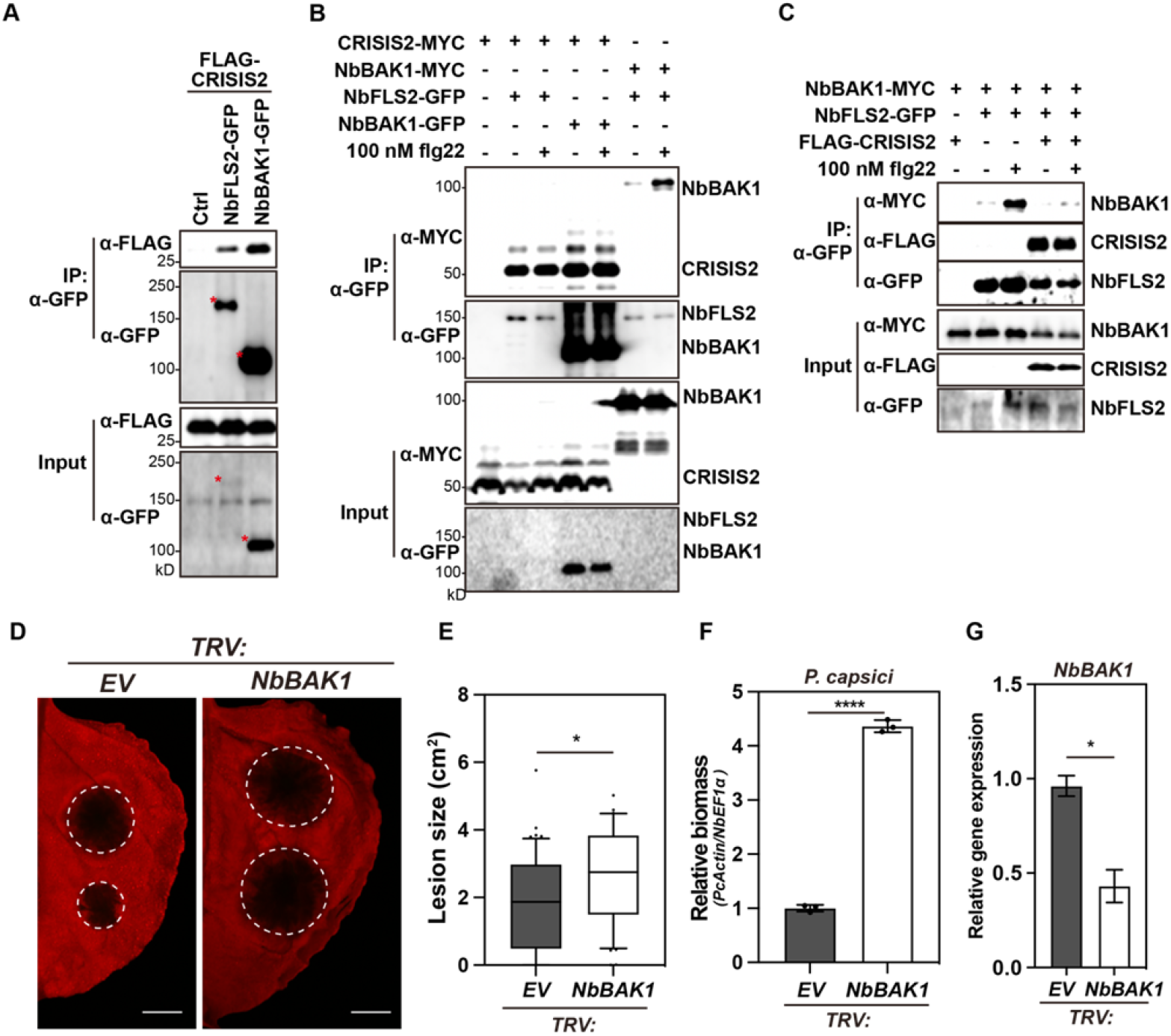
CRISIS2 inhibits the PAMP-induced PRR-BAK1 association. **A**, CRISIS2 associates with the NbFLS2 and NbBAK1 *in planta*. FLAG-CRISIS2 was co-expressed with Ctrl (empty vector), NbFLS2-GFP, or NbBAK1-GFP in *N. benthamiana* leaves. Proteins were extracted and subjected to immunoprecipitation (IP) with α-GFP agarose (IP: α-GFP) and immunoblotted with appropriate antibodies (α-FLAG, α-MYC and α-GFP). Input was collected from the same protein extracts before IP. **B**, The association of CRISIS2 and NbFLS2 or NbBAK1 is independent of flg22-induced PRR activation. CRISIS2-MYC and NbFLS2-GFP or NbBAK1 were co-infiltrated in *N. benthamiana* leaves by agro-infiltration. NbBAK1-MYC and NbFLS2-GFP were used as positive controls for flg22-induced association. At 48 hpi, water or 100 nM flg22 was treated for 15 min by infiltration. **C**, CRISIS2 interferes with the flg22-induced NbFLS2-NbBAK1 association. NbFLS2-GFP and NbBAK1-MYC were co-expressed with or without FLAG-CRISIS2 in *N. benthamiana* leaves, followed by 100 nM flg22 treatment. Proteins were extracted and subjected to immunoprecipitation (IP) with α-GFP agarose (IP: α-GFP) and immunoblotted with appropriate antibodies (α-FLAG, α-MYC and α-GFP). Input was collected from the same protein extracts before IP. **D**, Silencing of *NbBAK1* enhanced the virulence of *P. capsici*. *P. capsici* zoospores (4 × 10^4^ zoospores/ml) were drop-inoculated onto *EV-* or *NbBAK1-*silenced *N. benthamiana* leaves at 2 weeks after VIGS. The photographs were taken at 2 dpi. Bar = 1 cm. **E**, Lesion size of (D). The data are represented as 10-90% box and whisker (n = 40). Outliers are plotted as black dot and medians are the black lines in the boxes. The significant difference compared with *TRV:EV* was determined by unpaired t-test. *, p < 0.05; ***, p < 0.001. **F**, Biomass of *P. capsici* in (D). At 2 days after *P. capsici* inoculation, leaf disks around the inoculated site were harvested. The total genomic DNA was extracted and subjected to qPCR analysis. The biomass of *P. capsici* was determined by measuring *PcActin* gene normalized to *NbEF1⍺* gene. The significant difference compared with *TRV:EV* was determined by unpaired t-test. ****, p < 0.0001. **G**, Transcript accumulation measured by qRT-PCR to confirm the silencing efficiency of *NbBAK1*. The transcript accumulation of *NbBAK1* was measured at 2 weeks after VIGS. Data are represented as mean ± SD. Significant differences were determined by unpaired t-tests. *, p < 0.05. **J**, *P. capsici*-induced defense gene expression was reduced in the *fls2* mutant. Col-0 and *fls2* plants were inoculated with *P. capsici* zoospores for 1, 2, and 3 days. The expression of *FRK1* was examined using qRT-PCR analysis. Data are normalized to Col-0 1dpi and represented as mean ± SD. The significant difference was determined by two-way ANOVA (Sidak’s multiple comparisons test). *, p < 0.05; **, p < 0.01; ***, p < 0.001.

### Screening for CRISIS2 interactors revealed candidate PRRs involved in *P. capsici* detection

BAK1 functions as a regulatory hub of leucine-rich repeat receptor-like kinase (LRR-RLK) protein in immunity and has a general regulatory role in PM-associated receptor complexes (Couto and Zipfel, 2016). Considering that CRISIS2 suppressed multiple PTI responses against both flg22 and elf18, we hypothesized that CRISIS2 constitutively associates with NbBAK1 and consequently inhibits the interaction between BAK1 and unknown PRRs in response to *P. capsici*. To identify additional putative PRRs that are directly or indirectly targeted by CRISIS2, we performed *in planta* protein-protein interaction screening by transient expression of *GFP-CRISIS2* or *GFP* in *N. benthamiana* followed by anti-GFP co-IP and liquid chromatography tandem mass spectrometry (LC-MS/MS). This proteomic approach identified 339 putative host protein interactors specific for *GFP-CRISIS2* after subtraction of overlapping candidates from the GFP control (Supplemental Figure S11). Interestingly, NbPMA3 and Nb14-3-3 were validated as CRISIS2 interactors by this approach (Figure 5F). Moreover, additional PMAs were identified as interactors with high scores (Supplemental Figure S10B). These results indicated that the putative interactome revealed by our screen is likely to contain biologically relevant host target proteins.

We identified 3 putative receptor-like kinases (RLKs) including L-type lectin Receptor-Like Kinase (LecRLK), NbRLK1 and NbRLK5 as possible interactors of CRISIS2 (Supplemental Figure S10B). Co-IP confirmed the interaction between CRISIS2 and these three RLKs in *N. benthamiana* (Figure 9A). In addition, the three RLKs also associated with NbBAK1 in *N. benthamiana* (Figure 9B). To determine the role of these identified RLKs in the virulence of *P. capsici*, we silenced LecRLK, RLK1, or RLK5 in *N. benthamiana* using VIGS and monitored *P. capsici* infection in the silenced plants. Surprisingly, a significantly enhanced virulence with increased disease lesion size was observed in all three RLK gene-silenced plants compared with EV control (Figure 9C and 9D). Moreover, a significantly increased biomass of *P. capsici* was observed in all three RLK-silenced plants (Figure 9E). These results imply that the newly identified RLKs including LecRLK, RLK1 and RLK5 play important roles in plant defense and are the major targets of the RxLR effector CRISIS2 to inhibit PTI response during *P. capsici* infection. Thus, LecRLK, RLK1 and RLK5 appear as strong candidate PRRs, likely involved in the early detection of *P. capsici*-associated molecular patterns.

**Figure 9.**
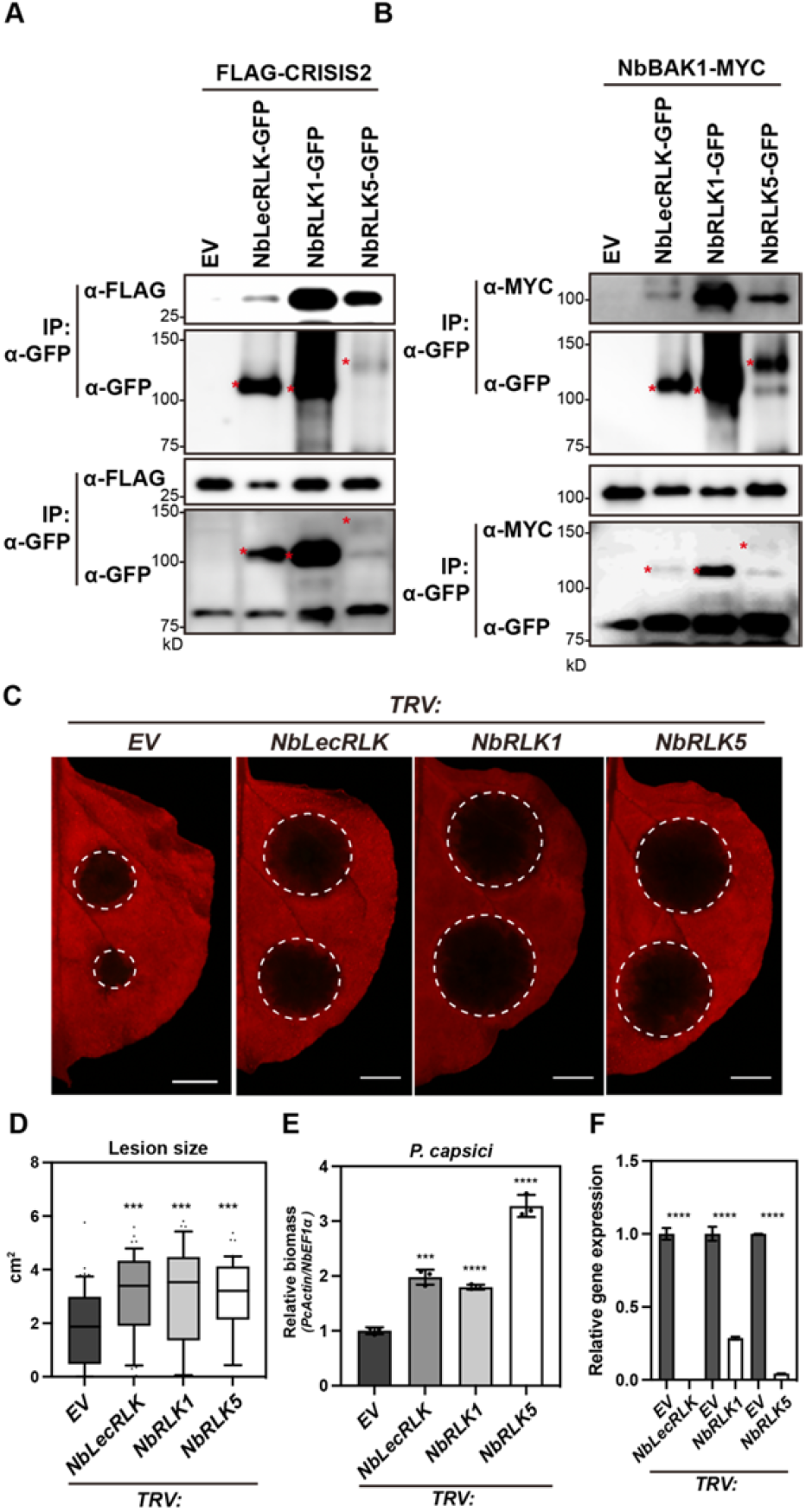
Host-interactor screen of CRISIS2 identifies putative PRRs in *N. benthamiana*. **A-B**, NbLecRLK, NbRLK1, and NbRLK5 interact with CRISIS2 and NbBAK1 *in planta*. FLAG-CRISIS2 (A) or NbBAK1-MYC (B) was transiently co-expressed with RLKs in *N. benthamiana* leaves by agro-infiltration. Proteins were extracted and subjected to immunoprecipitation (IP) with α-GFP agarose (IP: α-GFP) and immunoblotted with appropriate antibodies (α-FLAG, α-MYC and α-GFP). Input was collected from the same protein extracts before IP. **C**, Silencing of RLKs enhanced the virulence of *P. capsici*. Indicated genes were silenced in *N. benthamiana* by VIGS. *P. capsici* zoospores (4 × 10^4^ zoospores/ml) were drop-inoculated onto *N. benthamiana* leaves at 2 weeks after VIGS. The photographs were taken at 2 dpi. Bar = 1 cm. **D**, Lesion size of (C). The data are represented as 10-90% box and whisker (n ≥ 40). Outliers are plotted as black dot and medians are the black lines in the boxes. The significant difference compared with *TRV:EV* was determined by unpaired t-test. *, p < 0.05; ***, p < 0.001. **E**, Biomass of *P. capsici* in (C). At 2 days after *P. capsici* inoculation, leaf disks around the inoculated site were harvested. The total genomic DNA was extracted and subjected to qPCR analysis. The biomass of *P. capsici* was determined by measuring *PcActin* gene normalized to *NbEF1⍺* gene. The significant difference compared with *TRV:EV* was determined by unpaired t-test. ***, p < 0.001; ****, p < 0.0001. **F**, Transcript accumulation measured by qRT-PCR to confirm the silencing efficiency of indicated genes. The transcript level was measured at 2 weeks after VIGS. Data are represented as mean ± SD. Significant differences were determined by unpaired t-tests. ****, p < 0.0001.

## Discussion

Previously, we proposed PMA as the primary target of PM-associated CNLs to facilitate defense-associated cell death (Lee et al., 2022). Here, we hypothesized that pathogen effectors can also utilize the same machinery to promote pathogen virulence. PMAs are major proton pumps that build up an electrochemical gradient across the PM and PMAs are dynamically regulated during plant immune responses (Elmore and Coaker, 2011a). Moreover, several pathogenic microorganisms are known to modulate the PMA activity during infection (Elmore and Coaker, 2011b). Furthermore, distinct effectors from oomycetes and bacteria have the ability to suppress cell death induced by pathogen perception in plants (Bos *et al*., 2006; Dou et al., 2008). For example, of 169 *P. sojae* effectors tested, 127 effectors could consistently or partially suppress the cell death triggered by various elicitors of plant immune responses, while only 11 effectors triggered cell death (Yu et al., 2012), suggesting that they are likely recognized by the plant defense system or that they act as toxins in *N. benthamiana*. To overcome the experimental limitations due to the high number of effector candidates present in oomycete genomes, we designed multi-omics approaches and efficiently isolated 25 *P. capsici* RxLR effector candidates. Among them, 14 RxLR effectors have been confirmed to have a biological function referred to as cell death-inducing activity in the host plant *N. benthamiana*. Pharmacological screening using the irreversible PMA activator fusicoccin revealed that the cell death caused by 9 out of 14 effectors was affected by the PMA activator. However, only CRISIS2 physically and functionally associates with NbPMAs. It is widely accepted that irreversible activation of PMAs caused by fusicoccin results in disturbance of many secondary transport processes, such as sugar transport, nutrient uptake, and K^+^ uptake (Michelet and Boutry, 1995). Most likely, the imbalance across membrane potentials could be the cause of the suppression of cell death. Many pathogenic microorganisms may target PMA during infection (Elmore and Coaker, 2011). Often, the effective inhibition of proton pumping across the PM leads to depolarization and eventually cell death (Golstein and Kroemer, 2007). Moreover, hemibiotrophic and biotrophic fungal pathogens are known to cause extracellular alkalization in plants as an important feature of their lifestyles (Prusky et al., 2001; Prusky and Yakoby, 2003). Accordingly, great attention has been focused on identifying the pathogen or host components targeting PMA. To date, quite a few toxins from diverse pathogens have been reported to modulate PMA activity, including fusicoccin and NIP1 as activators and beticolin-1, fumonisin B1, bacterial lipopeptides, and tenuazonic acid as inhibitors (Bjørk et al., 2020). Several elicitors or PAMPs from common pathogens are also known to induce ion fluxes and rapid alkalization of the medium in cell suspension cultures, and it was suggested that those extracellular alkalizations may be due to the inhibition of PMA (Felix et al., 1999; Felix et al., 1993; Nürnberger et al., 1994). However, the molecular mechanism underlying the regulation of PMA remains elusive. Here, we demonstrated that a novel *P. capsici* RxLR effector, CRISIS2, modulates the activity of PMA by associating with the C-terminal regulatory domain of the enzyme to prevent the switch from an inactive to an active state of the enzyme.

Effectors can be recognized by plant immune receptors such as NLRs, leading to ETI, a stronger and efficient form of resistance that is frequently associated with HR cell death. Our previous study that PMA is a target of PM-associated CNLs to facilitate cell death led us to investigate whether CRISIS2-induced cell death is NLR-dependent. We confirmed that CRISIS2-induced cell death is not mediated by SGT1 and RAR1, the core regulators of NLR-mediated cell death (Azevedo et al., 2002; Boller and Felix, 2009), partially supporting our hypothesis that CRISIS2-induced cell death is unlikely due to recognition by NLR proteins but would rather support the necrotrophic stage of *P. capsici* through PMA inhibition.

We also have shown that CRISIS2 triggers cell death in a dose dependent manner based on transient expression using different binary vector systems and that only the CRISIS2 constructs inducing cell death could inhibit PMA activity in *N. benthamiana*. However, cell death-defective CRISIS2 still functions in promoting *P. capsici* virulence, suggesting that the ability to trigger cell death is not required for the suppression of defense responses in *N. benthamiana* when CRISIS2 is overexpressed. Whether CRISIS2-induced cell death indeed happens under natural conditions or is an artifact of elevated expression by transient overexpression is not clear. However, the pleiotropic physiological functions of CRISIS2 may be regulated with the *P. capsici* life cycle, biotrophic or necrotrophic phase, in the host plant.

Plant genomes encode a large family of RLKs that typically comprise extracellular receptor-like proteins with intracellular kinases. A large number of RLKs can specifically recognize PAMPs from pathogens and serve as PRRs in the PM (Böhm et al., 2014). A recent study reported that the *P. capsici* effector RxLR25 inhibits the phosphorylation of BIK1, a core component of FLS2-mediated signaling, to enhance virulence against pathogens (Liang et al., 2021). In this study, we demonstrated that CRISIS2 constitutively associated with BAK1, eventually disturbing the flg22-induced association between FLS2 and BAK1. We further obsevrved that CRISIS2 inhibits the elf18-induced PTI response. These results suggest that the major target of CRISIS2 is BAK1. Moreover, in later stage of this research, we identified NbLecRLK, an RLK with an extracellular legume-like lectin domain, NbRLK1, and NbRLK5 as a potential PRRs or PRR complex components likely involved in *P. capsici* detection (Hervé et al., 1999). In Arabidopsis plants, LecRKIX.1, LecRK-IX.2, and LecRK1.9, which belong to the lectin receptor kinase family, are known as positive regulators of resistance against two oomycete pathogens, *P. brassicae* and *P. capsici* (Wang et al., 2015, Bouwmeester et al., 2011). However, despite many efforts to elucidate the signaling pathways, the molecular mechanisms underlying LecRLK-dependent responses are still largely known. Perception of PAMPs triggers the rapid association of the corresponding receptors with other LRR-RLK and induced PRR-RLK complex formation is an important feature for transducing immune signal. It would be interesting to explore what signaling cue or PAMP enhances or modulate the association between LecRLK and BAK1 during *Phytophthora* invasion.

Taken together, we propose a working model for CRISIS2 regulation of plant immune components to support the *P. capsici* hemibiotrophic lifestyle in a simultaneous or time-dependent manner (Figure S12). At the very early phase of infection, CRISIS2 may inhibit pathogen recognition by interrupting ligand-induced receptor complex formation. In the late phase of infection, CRISIS2 eventually contribute to host cell death by inhibiting PMA activity. Our findings that one RxLR effector has multiple targets and modulates different layers of the plant defenses support novel mechanisms of pathogenicity and provide new insight into how hemibiotrophic pathogens have evolved molecular weapons to survive in line with the evolution of the plant immune response against pathogen infection.

## Methods

### Plant materials and growth condition

*N. benthamiana* plants were grown in a walk-in chamber under a 16-h day and 8-h night at 25°C. Four- to five-week-old plants were used in the *Agrobacterium*-mediated transient overexpression assay, and two-week-old plants were used in the virus-induced gene silencing (VIGS) assay.

### Identification and selection of *P. capsici* RXLR effector

Phyca11 scaffolds and proteins were obtained from the *P. capsici* sequencing consortium website (https://mycocosm.jgi.doe.gov/Phyca11/Phyca11.home.html.). To identify *P. capsici* RxLR effector candidates, two independent approaches were used. One data set was generated by prediction of RxLR effectors using effetR software from all open reading frames (ORFs) extracted from Phyca11 scaffolds (EMBOSS getORF) (Rice et al., 2000, Tabima and Grunwald, 2019). Among the predicted effectors, RxLRs without signal peptides were filtered out using signalP 5.0 (Armenteros et al., 2019). Another data set was generated by TGFam-Finder using Phyca11 scaffolds (Kim et al., 2020). InterProScan5 was used for domain identification by setting the target domain ID to PF16810 according to the Pfam protein family database (Jones et al., 2014). Resource proteins used in TGFam-Finder were putative RxLR effectors of 20 oomycete species, in which the candidate RxLR effectors were extracted from genome data using effectR and signalP5.0. The 20 oomycete genome data were obtained from the Oomycete Gene Order Browser (OGOB; Mcgowan et al., 2019). Among the putative RxLR effectors that overlapped in the two separate data sets, 25 candidates that were differentially expressed during the early infection stage and biotrophic phase in the microarray expression profiles of a previous study were selected (Jupe et al., 2013).

### DNA construct

The effector domain of 24 selected *P. capsici* RXLR effector candidates was synthesized artificially in pTwist cloning vectors (LNCbio, Korea). They were amplified and cloned into the potato virus X-based vector pICH31160 (pKW) by the ligation-independent cloning (LIC) method (Oh et al., 2010). CRISIS2 and CRISIS6 were amplified and inserted into pKW-3xFLAG-LIC and pCAMBIA2300-LIC by the LIC method and into the pK7WGF2 and pXVE-DC-6xmyc vectors by gateway cloning (Invitrogen, USA). The primers used for construction are provided in Table S2. NbPMA1 (Niben101Scf00593g01002.1), NbPMA3 (Niben101Scf07395g00031.1), NbPMA4 (Niben101Scf03979g02010.1), NbFLS2(Niben101Scf01785g10011.1), NbAUX1 (Niben101Scf02269g03006.1), NbCNGC4(Niben101Scf04528g09004.1), Nb14-3-3(Niben101Scf02537g00004.1), NbBAK1 (Niben101Scf11779g01002.1), NbLecRLK (Niben101Scf07589g01014.1), NbRLK1 (Niben101Scf18639g00026.1), NbRLK5 (Niben101Scf00742g01037.1)

### Transient expression in *N. benthamiana*

For *in planta* transient overexpression, *Agrobacterium tumefaciens* strain GV3101 containing the desired expression vector was grown overnight at 28°C in LB media with appropriate antibiotics. The cells were collected by centrifugation at 3,000 rpm for 10 min and resuspended in infiltration buffer (10 mM MES (1-[N-morpholino] ethanesulfonic acid), 10 mM MgCl_2_ and 150 μM acetosyringone, pH 5.6). The resuspended cells were adjusted at OD_600_ = 0.5. For co-infiltration, an equal volume of each cell resuspension was mixed. In all transient overexpression experiments, a p19 silencing suppressor was included, and pressure was infiltrated using a needleless syringe.

### Virus-induced gene silencing (VIGS) in *N. benthamiana*

VIGS was performed by following Liu et al. (Liu et al., 2002). The suspensions of *A. tumefaciens* carrying TRV1 and TRV2 with the target gene fragment were mixed at a 1:1 ratio in infiltration buffer to a final O. D_600_ of 1.5 and infiltrated into two leaves of 2-week-old *N. benthamiana*. Three weeks later, the upper leaves were used for further experiments. To confirm the silencing efficiency, the transcript levels of the genes were validated by quantitative RT-PCR (qRT-PCR). Total RNA was extracted from silenced leaves using TRIzol reagent (MRC, USA), and cDNA was synthesized using SuperScript II Reverse Transcriptase (Invitrogen, USA). qRT-PCR was performed using a CFX96 Touch Real-Time PCR Detection System (Bio-Rad, USA) with ExcelTaq™ 2X Q-PCR Master Mix (SYBR, ROX; SMObio). The transcript level was normalized to that of the internal standard *elongation factor-1a* of *N. benthamiana* (*NbEF-1a*). The primers used for qRT-PCR are provided in Table S2.

### Host-induced gene silencing (HIGS) in *P. capsici*

HIGS was performed by using modified VIGS system. *CRISIS2*-specific DNA fragment was amplified from *P. capsici* cDNA and inserted into TRV2-LIC vector. The suspensions of *A. tumefaciens* carrying TRV1 and TRV2:CRISIS2 were mixed at a 1:1 ration in infiltration buffer to a final O. D_600_ of 0.15 and infiltrated into two leaves of 3-week-old *N. benthamiana*. After 10-14 days, the upper leaves were subjected to *P. capsici* inoculation.

### *P. capsici* culture conditions and inoculation assay and biomass measurement

*P. capsici* strain 40476 was routinely maintained on V8 agar medium at 23°C in the dark. For inoculation on *N. benthamiana*, mycelium was grown in V8 agar medium at 23°C in the dark for one week, and then the mycelia were scraped and incubated under white light for 12 h for sporulation. To release zoospores, sporangia were collected in distilled water and incubated at room temperature for 1 h. Four hundred zoospores were inoculated by drop inoculation on *N. benthamiana* and incubated in a growth chamber at 25°C for 2 days. The lesion size was measured at 2 dpi using ImageJ software. To measure the biomass of *P. capsici*, leaf disks around the inoculated site were harvested, and total genomic DNA was extracted. Biomass was determined by qPCR measuring the *PcActin* gene normalized to *NbEF1⍺*.

### ROS accumulation

Four-week-old *N. benthamiana* plants were infiltrated with Agrobacterium harboring *GFP, GFP-CRISIS2* or *GFP-SRISIS6*, and 12 leaf discs of 0.25 cm^2^ from 2 independent leaves were harvested and incubated overnight in a 96-well plate with 200 μL of distilled water (D. W) to eliminate the wounding stress. D. W was replaced by 100 μL of reaction solution including 50 μM luminol and 10 μg/mL horseradish peroxidase (Sigma-Aldrich) and 100 nM flg22 or elf 18. The measurement was performed immediately after adding the reaction solution with a 2-min interval and over a period of 60 min in a luminometer (Perkin-Elmer 2030 Multilabel Reader, Victor X3, USA). The measured value for ROS production from 24 leaf discs per treatment was indicated as the means of relative light units and repeated at least three times.

### MAPK assay

The MAPK assay was performed as described previously (Mang et al., 2017). Briefly, 4-week-old *N. benthamiana* were infiltrated with agrobacterium harboring *GFP, GFP-CRISIS2* or *GFP-SRISIS6*, and at least 8 leaf discs of 0.25 cm^2^ from 2 independent leaves were harvested and incubated for at least 6 hr in an 8-well plate with 1 mL of D. W to eliminate the wounding stress. D. W was replaced by 1 mL of D. W containing 100 μM flg22 and harvested at various time points. Samples were ground in 10 μl/one leaf disk of extraction buffer (150 mM NaCl, 50 mM Tris-HCl, pH 7.5, 5 mM EDTA, 1% Triton X-100, 2 mM Na_3_VO_4_, 2 mM NaF, 1 mM DTT, and 1:100 complete protease inhibitor cocktail of Sigma-Aldrich). The supernatant was collected after centrifugation at 12,000 rpm for 10 min at 4°C, and protein samples with 5X SDS buffer were loaded on 8% SDS-PAGE gels to detect pMPK3, pMPK6 and pMPK4 by immunoblotting with an α-pERK1/2 antibody (Cell Sinaling; no. 9101, USA).

### Co-IP and immunoblot assay

*Agrobacterium*-infiltrated *N. benthamiana* leaves were sampled at 36-48 hpi for co-IP or western blot. The GFP-tagged proteins were immunoprecipitated with 15 μl of anti-GFP agarose beads (MBL) in 700 μl co-IP buffer (10% glycerol, 50 mM Tris-HCl (pH 7.5), 2 mM EDTA, 150 mM NaCl, 10 mM DTT, 2% polyvinylpolypyrrolidone, 0.25% Triton X-100, 1:100 complete protease inhibitor cocktail (Roche, USA). Input was prepared by 50 μl aliquots of samples in co-IP buffer before adding anti-GFP agarose beads. The co-IP samples were gently rotated overnight at 4 °C. The beads were collected and washed six times with washing buffer (500 mM NaCl, 25 mM Tris-HCl (pH 7.5), 1 mM EDTA and 0.15% NP-40). The samples were eluted in 15 μl loading buffer and subjected to subsequent immunoblotting assays with appropriate antibodies.

### Visualization and quantification of cell death

The degree of cell death was measured by chlorophyll fluorescence using a closed FluorCam (Photon Systems Instruments, CZ) and quantified by FluorCam 7.0 software. Leaves infiltrated with agrobacterium with each plasmid were detached and exposed to a superpulse in a closed chamber, and minimum fluorescence (F0), maximum fluorescence (Fm), and maximum quantum yield of photosystem II (Fv/Fm) parameters were determined using the default Fv/Fm protocol.

### Confocal Microscopy

pH measurement was performed as previously described (Lee et al., 2021). Briefly, transgenic *N. benthamiana* transformed with PM-APO, a ratiometric pHluorin sensor for apoplast, was infiltrated with Agrobacterium harboring EV- or FLAG-CRISIS2. Confocal microscopic observation and quantification of fluorescence signals were performed as described, with modifications. Observations were performed with a Leica SP8 X microscope using a 20X water objective with the same WLL laser at 40% 476 nm and at 20% 496 nm output. Emission was detected at 505 and 550 nm, with the pinhole set to 1 airy unit. Subcellular localization of CRISISs was observed using a confocal microscope (Leica SP8 X, Germany). The fluorescence of eGFP and mStrawberry was detected under 488 nm excitation and 500-550 emission or 574 nm excitation and 610-650 nm emission, respectively. For plasmolysis, the leaf disk was incubated in 1 M NaCl solution for 15 min. Images were processed using LAS X software.

### Yeast two hybrid assay

CRISIS2 or Nb14-3-3 was fused with the Gal4 DNA binding domain in pGBKT7 (Clontech, PT3248-5, USA) and inserted into the *Saccharomyces cerevisiae* Y2HGold strain (Clontech, USA) under selection with SD/-Trp. Cytosolic domain fragments of NbPMA3 (F1, F2 or F3) were fused with the Gal4 activation domain in pGADT7 (Clontech, PT3249-5, USA) and inserted into the Y187 strain under selection with SD/-Leu. Lam-pGBKT7 was used as a negative control for bait against each pGADT7 construct. For the positive control, a combination of Nb14-3-3-pGBKT7 and NbPMA3(F3)-pGADT7 was used. After mating, the co-transformants containing pGBKT7 and pGADT7 were selected on SD/-Trp/-Leu and SD/-Trp/-Leu/-His media. The plates were incubated at 30°C for 7 days.

### Statistical Analysis

Graphs generation and statistical test indicated at figure legends were performed with PRISM 9 (graphPad). Error bars represent standard deviation or standard error of mean. Student’s t-tests, one-way ANOVA followed by Tukey’s test, or two-way ANOVA followed by Sidak’s multiple comparisons test was used.

## Acknowledgments

We thank Yubin Lee for preparing experiments. This work was supported by a National Research Foundation of Korea (NRF) grant funded by the Korean government (MSIT) (No. 2018R1A5A1023599 [SRC] and 2021R1A2B5B03001613) to D.C., The authors have no conflicts of interest to declare.

## Author contributions

D.C. conceived the project; H.M., Y.-E.S., designed the experiments; H.M., Y.-E.S., H.-Y.L., H.K., H.J., X.Y., S.P., M.-S.K., C.S. performed the experiments; H.M., Y.-E.S., C.S., and D.C. wrote the article with input from the other authors.

## Competing interests

The authors declare no competing interests.

## Parsed Citations

Avrova, A.O., Boevink, P.C., Young, V., Grenville-Briggs, L.J., Van West, P., Birch, P.R., and Whisson, S.C. (2008). Anovel Phytophthora infestans haustorium-specific membrane protein is required for infection of potato. Cell. microbiol. 10: 2271–2284.

Azevedo, C., Sadanandom, A., Kitagawa, K., Freialdenhoven, A., Shirasu, K., and Schulze-Lefert, P. (2002). The RAR1 interactor SGT1, an essential component of R gene-triggered disease resistance. Science 295: 2073–2076.

Baunsgaard, L., Fuglsang, A.T., Jahn, T., Korthout, H., de Boer, A.H., and Palmgren, M.G. (1998). The 14-3-3 proteins associate with the plant plasma membrane H (+)-ATPase to generate a fusicoccin binding complex and a fusicoccin responsive system. Plant J. 13: 661–671.

Bjørk, P.K., Rasmussen, S.A., Gjetting, S.K., Havshøi, N.W., Petersen, T.I., Ipsen, J.Ø., Larsen, T.O., and Fuglsang, A.T. (2020). Tenuazonic acid from Stemphylium loti inhibits the plant plasma membrane H+-ATPase by a mechanism involving the C-terminal regulatory domain. New Phytol. 226: 770–784.

Block, A., Li, G., Fu, Z.Q., and Alfano, J.R. (2008). Phytopathogen type III effector weaponry and their plant targets. Curr. Opin. Plant Biol. 11: 396–403.

Boller, T., and Felix, G. (2009). Arenaissance of elicitors: perception of microbe-associated molecular patterns and danger signals by pattern-recognition receptors. Annu. Rev. Plant Biol. 60: 379–406.

Boller, T., and He, S.Y. (2009). Innate immunity in plants: an arms race between pattern recognition receptors in plants and effectors in microbial pathogens. Science 324: 742–744.

Bos, J.I., Armstrong, M.R., Gilroy, E.M., Boevink, P.C., Hein, I., Taylor, R.M., Zhendong, T., Engelhardt, S., Vetukuri, R.R., and Harrower, B. (2010). Phytophthora infestans effector AVR3a is essential for virulence and manipulates plant immunity by stabilizing host E3 ligase CMPG1. Proc. Natl. Acad. Sci. 107: 9909–9914.

Bos, J.I., Kanneganti, T.D., Young, C., Cakir, C., Huitema, E., Win, J., Armstrong, M.R., Birch, P.R., and Kamoun, S. (2006). The C-terminal half of Phytophthora infestans RXLR effector AVR3a is sufficient to trigger R3a-mediated hypersensitivity and suppress INF1-induced cell death in Nicotiana benthamiana. Plant J. 48: 165–176.

Botër,.M., Amigues, B., Peart, J., Breuer, C., Kadota, Y., Casais, C., Moore, G., Kleanthous, C., Ochsenbein, F., and Shirasu, K. (2007). Structural and functional analysis of SGT1 reveals that its interaction with HSP90 is required for the accumulation of Rx, an R protein involved in plant immunity. Plant Cell 19: 3791–3804.

Bozkurt, T.O., Schornack, S., Win, J., Shindo, T., Ilyas, M., Oliva, R., Cano, L.M., Jones, A.M., Huitema, E., and van der Hoorn, R.A. (2011). Phytophthora infestans effector AVRblb2 prevents secretion of a plant immune protease at the haustorial interface. Proc. Natl. Acad. Sci. 108: 20832–20837.

Chaparro-Garcia, A., Wilkinson, R.C., Gimenez-Ibanez, S., Findlay, K., Coffey, M.D., Zipfel, C., Rathjen, J.P., Kamoun, S., and Schornack, S. (2011). The receptor-like kinase SERK3/BAK1 is required for basal resistance against the late blight pathogen Phytophthora infestans in Nicotiana benthamiana. PloS one 6: e16608.

Chen, S., Yin, C., Qiang, S., Zhou, F., and Dai, X. (2010). Chloroplastic oxidative burst induced by tenuazonic acid, a natural photosynthesis inhibitor, triggers cell necrosis in Eupatorium adenophorum Spreng. Biochim. Biophys. Acta-Bioenerg. 1797: 391–405.

Chinchilla, D., Zipfel, C., Robatzek, S., Kemmerling, B., Nürnberger, T., Jones, J.D., Felix, G., and Boller, T. (2007). Aflagellin-induced complex of the receptor FLS2 and BAK1 initiates plant defence. Nature 448: 497–500.

Couto, D., and Zipfel, C. (2016). Regulation of pattern recognition receptor signalling in plants. Nat. Rev. Immunol. 16: 537–552.

Dou, D., Kale, S.D., Wang, X., Jiang, R.H., Bruce, N.A., Arredondo, F.D., Zhang, X., and Tyler, B.M. (2008). RXLR-mediated entry of Phytophthora sojae effector Avr1b into soybean cells does not require pathogen-encoded machinery. Plant Cell 20: 1930–1947.

Dou, D., Kale, S.D., Wang, X., Chen, Y., Wang, Q., Wang, X., Jiang, R.H., Arredondo, F.D., Anderson, R.G., and Thakur, P.B. (2008). Conserved C-terminal motifs required for avirulence and suppression of cell death by Phytophthora sojae effector Avr1b. Plant Cell 20: 1118–1133.

Elmore, J.M., and Coaker, G. (2011). The role of the plasma membrane H+-ATPase in plant–microbe interactions. Mol. Plant 4: 416–427.

Felix, G., Duran, J.D., Volko, S., and Boller, T. (1999). Plants have a sensitive perception system for the most conserved domain of bacterial flagellin. Plant J. 18: 265–276.

Felix, G., Regenass, M., and Boller, T. (1993). Specific perception of subnanomolar concentrations of chitin fragments by tomato cells: induction of extracellular alkalinization, changes in protein phosphorylation, and establishment of a refractory state. Plant J. 4: 307–316.

Felle, H.H., Herrmann, A., Hanstein, S., Hückelhoven, R., and Kogel, K.-H. (2004). Apoplastic pH signaling in barley leaves attacked by the powdery mildew fungus Blumeria graminis f. sp. hordei. Mol. Plant-Microbe Interact. 17: 118–123.

Heese, A., Hann, D.R., Gimenez-Ibanez, S., Jones, A.M., He, K., Li, J., Schroeder, JI., Peck, S.C., and Rathjen, J.P. (2007). The receptor-like kinase SERK3/BAK1 is a central regulator of innate immunity in plants. Proc. Natl. Acad. Sci. 104: 12217–12222.

Jupe, F., Witek, K., Verweij, W., Śliwka, J., Pritchard, L., Etherington, G.J., Maclean, D., Cock, P.J., Leggett, R.M., and Bryan, G.J. (2013). Resistance gene enrichment sequencing (R en S eq) enables reannotation of the NB-LRR gene family from sequenced plant genomes and rapid mapping of resistance loci in segregating populations. Plant J. 76: 530–544.

Kadota, Y., Sklenar, J., Derbyshire, P., Stransfeld, L., Asai, S., Ntoukakis, V., Jones, J.D., Shirasu, K., Menke, F., and Jones, A. (2014). Direct regulation of the NADPH oxidase RBOHD by the PRR-associated kinase BIK1 during plant immunity. Mol. Cell 54: 43–55.

Keinath, N.F., Kierszniowska, S., Lorek, J., Bourdais, G., Kessler, S.A., Shimosato-Asano, H., Grossniklaus, U., Schulze, W.X., Robatzek, S., and Panstruga, R. (2010). PAMP (pathogen-associated molecular pattern)-induced changes in plasma membrane compartmentalization reveal novel components of plant immunity. J. Biol. Chem. 285: 39140–39149.

Kim, S., Cheong, K., Park, J., Kim, M.S., Kim, J., Seo, M.K., Chae, G.Y., Jang, M.J., Mang, H., and Kwon, S.H. (2020). TGFam-Finder: a novel solution for target-gene family annotation in plants. New Phytol. 227: 1568–1581.

Koeck, M., Hardham, A.R., and Dodds, P.N. (2011). The role of effectors of biotrophic and hemibiotrophic fungi in infection. Cell. Microbiol. 13:1849–1857.

Kühlbrandt, W. (2004). Biology, structure and mechanism of P-type ATPases. Nat. Rev. Mol. Cell Biol. 5: 282–295.

Kunze, G., Zipfel, C., Robatzek, S., Niehaus, K., Boller, T., and Felix, G. (2004). The N terminus of bacterial elongation factor Tu elicits innate immunity in Arabidopsis plants. Plant Cell 16: 3496–3507.

Lacomme, C., and Chapman, S. (2008). Use of Potato Virus X (PVX)–Based Vectors for Gene Expression and Virus-Induced Gene Silencing (VIGS). Curr.Protoc. Microbiol. 8: 16I–1.

Lee, H.Y., Seo, Y.E., Lee, J.H., Lee, S.E., Oh, S., Kim, J., Jung, S., Kim, H., Park, H., and Kim, S. (2022). Plasma membrane-localized plant immune receptor targets H+-ATPase for membrane depolarization to regulate cell death. New Phytol. 233: 934–947.

Li, Q., Ai, G., Shen, D., Zou, F., Wang, J., Bai, T., Chen, Y., Li, S., Zhang, M., and Jing, M. (2019). APhytophthora capsici effector targets ACD11 binding partners that regulate ROS-mediated defense response in Arabidopsis. Mol. Plant 12: 565–581.

Liang, X., Bao, Y., Zhang, M., Du, D., Rao, S., Li, Y., Wang, X., Xu, G., Zhou, Z., and Shen, D. (2021). APhytophthora capsici RXLR effector targets and inhibits the central immune kinases to suppress plant immunity. New Phytol. 232: 264–278.

Maekawa, T., Kufer, T.A., and Schulze-Lefert, P. (2011). NLR functions in plant and animal immune systems: so far and yet so close. Nat. Immunol. 12: 817–826.

Mang, H., Feng, B., Hu, Z., Boisson-Dernier, A., Franck, C.M., Meng, X., Huang, Y., Zhou, J., Xu, G., and Wang, T. (2017). Differential regulation of two-tiered plant immunity and sexual reproduction by ANXUR receptor-like kinases. Plant Cell 29: 3140–3156.

Melech-Bonfil, S., and Sessa, G. (2010). Tomato MAPKKKε is a positive regulator of cell-death signaling networks associated with plant immunity. Plant J. 64: 379–391.

Monaghan, J., and Zipfel, C. (2012). Plant pattern recognition receptor complexes at the plasma membrane. Curr. Opin. Plant Biol. 15: 349–357.

Moon, J.Y., Lee, J.H., Oh, C.S., Kang, H.G., and Park, J.M. (2016). Endoplasmic reticulum stress responses function in the HRT- mediated hypersensitive response in Nicotiana benthamiana. Mol. Plant Pathol. 17: 1382–1397.

Mucyn, T.S., Clemente, A., Andriotis, V.M., Balmuth, A.L., Oldroyd, G.E., Staskawicz, B.J., and Rathjen, J.P. (2006). The tomato NBARC-LRR protein Prf interacts with Pto kinase in vivo to regulate specific plant immunity. Plant Cell 18: 2792–2806.

Nürnberger, T., Nennstiel, D., Jabs, T., Sacks, W.R., Hahlbrock, K., and Scheel, D. (1994). High affinity binding of a fungal oligopeptide elicitor to parsley plasma membranes triggers multiple defense responses. Cell 78: 449–460.

Oh, S.-K., Kim, S.-B., Yeom, S.-I., Lee, H.-A., and Choi, D. (2010). Positive-selection and ligation-independent cloning vectors for large scale in planta expression for plant functional genomics. Mol. Cells 30: 557–562.

Oh, S.-K., Kim, H., and Choi, D. (2014). Rpi-blb2-mediated late blight resistance in Nicotiana benthamiana requires SGT1 and salicylic acid-mediated signaling but not RAR1 or HSP90. FEBS Lett. 588: 1109–1115.

Rice, P., Longden, I., and Bleasby, A. (2000). EMBOSS: the European molecular biology open software suite. Trends Genet. 16: 276–277.

Shan, L., He, P., Li, J., Heese, A., Peck, S.C., Nürnberger, T., Martin, G.B., and Sheen, J. (2008). Bacterial effectors target the common signaling partner BAK1 to disrupt multiple MAMP receptor-signaling complexes and impede plant immunity. Cell Host Microbe 4: 17–27.

Tyler, B.M., Tripathy, S., Zhang, X., Dehal, P., Jiang, R.H., Aerts, A., Arredondo, F.D., Baxter, L., Bensasson, D., and Beynon, J.L. (2006). Phytophthora genome sequences uncover evolutionary origins and mechanisms of pathogenesis. Science 313: 1261–1266.

Vera-Estrella, R., Barkla, B.J., Higgins, V.J., and Blumwald, E. (1994). Plant defense response to fungal pathogens (activation of host-plasma membrane H+-ATPase by elicitor-induced enzyme dephosphorylation). Plant Physiol. 104: 209–215.

Wang, Y., Bouwmeester, K., Beseh, P., Shan, W., and Govers, F. (2014). Phenotypic analyses of Arabidopsis T-DNAinsertion lines and expression profiling reveal that multiple L-type lectin receptor kinases are involved in plant immunity. Mol. Plant-Microbe Interact. 27: 1390–1402.

Wang, Y., Cordewener, J.H., America, A.H., Shan, W., Bouwmeester, K., and Govers, F. (2015). Arabidopsis lectin receptor kinases LecRK-IX. 1 and LecRK-IX. 2 are functional analogs in regulating Phytophthora resistance and plant cell death. Mol. Plant-Microbe Interact. 28: 1032–1048.

Whisson, S.C., Boevink, P.C., Moleleki, L., Avrova, A.O., Morales, J.G., Gilroy, E.M., Armstrong, M.R., Grouffaud, S., Van West, P., and Chapman, S. (2007). Atranslocation signal for delivery of oomycete effector proteins into host plant cells. Nature 450: 115–118.

Xiang, T., Zong, N., Zou, Y., Wu, Y., Zhang, J., Xing, W., Li, Y., Tang, X., Zhu, L., and Chai, J. (2008). Pseudomonas syringae effector AvrPto blocks innate immunity by targeting receptor kinases. Curr. Biol. 18: 74–80.

Yang, K.-Y., Liu, Y., and Zhang, S. (2001). Activation of a mitogen-activated protein kinase pathway is involved in disease resistance in tobacco. Proc. Natl. Acad. Sci. 98: 741–746.

Yu, H., Yan, J., Du, X., and Hua, J. (2018). Overlapping and differential roles of plasma membrane calcium ATPases in Arabidopsis growth and environmental responses. J. Exp. Bot. 69: 2693–2703.

Yu, X., Tang, J., Wang, Q., Ye, W., Tao, K., Duan, S., Lu, C., Yang, X., Dong, S., and Zheng, X. (2012). The RxLR effector Avh241 from Phytophthora sojae requires plasma membrane localization to induce plant cell death. New Phytol. 196: 247–260.

Zhou, F., Andersen, C.H., Burhenne, K., Fischer, P.H., Collinge, D.B., and Thordal-Christensen, H. (2000). Proton extrusion is an essential signalling component in the HR of epidermal single cells in the barley–powdery mildew interaction. Plant J. 23: 245–254.

Zhu, L., Zhu, J., Liu, Z., Wang, Z., Zhou, C., and Wang, H. (2017). Host-induced gene silencing of rice blast fungus Magnaporthe oryzae pathogenicity genes mediated by the brome mosaic virus. Genes 8: 241.

